# HMGA2 Expression in Pancreatic Ductal Adenocarcinoma Controls Tumor Microenvironment and Immunosuppression via IL1α

**DOI:** 10.64898/2026.10.02.754697

**Authors:** Naomi Yamamoto, Stephanie Dobersch, Tess M. Robertson, Jae Hyung Sim, Adrianne Wallace-Povirk, Nayvin W. Chew, Alexander E. Orellana, Joana Aggrey-Fynn, Liberalis Debraj Boila, Aidan Schutter, Ian M. Loveless, Gun Ho Jang, Shayan Avanessian, Gregory L. Beatty, Sunil R. Hingorani, Camilla Rydberg Millrud, Nina G. Steele, Debora Barbosa Vendramini-Costa, Ralph Francescone, Howard C. Crawford, Faiyaz Notta, Kevin C. Barry, Sita Kugel

**Author notes:** Corresponding author. Mail: 1100 Fairview Avenue N, Mail Stop C1-015, Seattle, WA 98109.

## Abstract

The basal-like subtype of pancreatic ductal adenocarcinoma (PDAC) is associated with poor outcomes and increased chemoresistance. Expression of high mobility group AT-hook 2 (HMGA2), a chromatin architectural protein, marks basal PDAC cells and associates with specific features of the tumor microenvironment (TME), including more immunosuppressive cancer associated fibroblasts (CAFs) and fewer cytotoxic CD8^+^ T cells. Expression of HMGA2 by tumor cells leads to upregulation of the immunosuppressive markers interleukin 6 (IL6) and leucine-rich repeat containing protein 15 (LRRC15) in associated fibroblasts. Mechanistically, HMGA2 in the tumor cell binds the interleukin 1 alpha (IL1α) promoter and upregulates transcription, increasing IL1α in the TME, in turn inducing immunosuppressive fibroblast polarization, increasing regulatory T cells, and decreasing CD8^+^ T cells. Blockade of the IL1α signaling axis disrupts this tumor-driven microenvironment phenotype and combines with inhibitors of oncogenic KRAS to improve tumor control.

**Statement of Significance:** The tumor microenvironment of pancreatic ductal adenocarcinoma drives immunosuppression, but causal relationships between PDAC subtypes and the TME have not been established. We report direct regulation of IL1α signaling by a basal PDAC-specific factor that controls fibroblast polarization and drives immunosuppression.

## INTRODUCTION

Pancreatic ductal adenocarcinoma (PDAC) is an epithelial tumor arising from neoplastic ducts and acinar-to-ductal metaplasia in the exocrine pancreas. While incidence of most common cancers is decreasing, the incidence of PDAC is on the rise in the United States and internationally^1,2^. Most cases of PDAC are diagnosed after the tumor has already metastasized, precluding curative resection and leading to a dismal five-year survival rate below 10%^1,2^. The mainstay of treatment for metastatic PDAC remains combination chemotherapy. While transformative in other cancers, immunotherapy has yet to impact PDAC treatment^3^. Taken together, these factors have resulted in universally poor outcomes, especially in economically disadvantaged and under-resourced settings^4,5^.

Multiple landmark studies have identified two main transcriptional subsets: “classical,” which is associated with more favorable outcomes and expression of normal pancreatic lineage genes, and “basal-like,” which is associated with worse outcomes and expression of mesenchymal genes and squamous signatures^6–11^. Understanding the origin, maintenance, and therapeutic vulnerabilities of these subtypes may lead to full implementation of cell-state subtyping in the clinic to drive therapeutic benefit. While classical subtype PDAC appears to respond better to chemotherapy^9,12,13^, basal tumors have been reported to be more responsive to KRAS inhibition^14^, EGFR inhibition^15^, CDK7 inhibition^16^, and protein synthesis inhibition^17^, among others. PDAC is notoriously refractory to current immunotherapy regimens, except for patients with germline or somatic mutations to mismatch repair machinery^18–20^.

PDAC resistance to immunotherapy has been attributed to its complex tumor microenvironment (TME), where desmoplastic fibroblasts and myeloid-derived suppressor cells predominate the landscape^21^. PDAC also demonstrates a relative paucity of CD8^+^ T cells compared to other solid malignancies where immunotherapies have entered clinical practice^22–24^. The immunosuppression seen in PDAC is due, at least in part, to the desmoplastic response during pancreatic tumorigenesis. Many types of cancer-associated fibroblasts (CAFs) have been well-characterized using patient samples and mouse models of PDAC, including inflammatory CAFs (iCAFs), myofibroblastic CAFs (myCAFs), and antigen-presenting CAFs (apCAFs)^25–30^. Additional markers for PDAC CAFs have also been described, including NetG1^31^, MHCII^28^, and LRRC15^32^. Previous studies have linked the basal transcriptional subtype to a homogeneous TME^8^, paucity of CD4^+^ and CD8^+^ T cells^8,13,33^, and associations with both immunosuppressive fibroblast activation protein (FAP) high CAFs^13,33,34^ and various subtypes of myCAFs^35,36^. However, the causal link between PDAC tumor transcriptional subtype and the functionality of the associated CAFs and resulting TME remains incompletely understood^37,38^.

We have previously described expression of the oncofetal chromatin-associated protein high-mobility group protein AT-hook 2 (HMGA2) as a marker of the basal subtype and chemoresistance in patient samples^13^, and found that expression is sufficient to drive the basal state *in vitro* and *in vivo*^17^. Here, we demonstrate that HMGA2 expression in tumor cells regulates IL1α expression, which polarizes fibroblasts toward an immunomodulatory IL6^+^ LRRC15^+^ CAF phenotype and ultimately drives immunosuppression in the PDAC TME.

## RESULTS

### Basal HMGA2^+^ tumor epithelial cells associate with an immunosuppressive fibroblast phenotype

We previously demonstrated an association between high tumoral expression of the basal marker *HMGA2* and high expression of *FAP* in CAFs using both multiplex immunohistochemistry (IHC) and single-cell RNA sequencing (scRNAseq) in patient samples^13^. Using data from spatial transcriptomics studies in end-stage metastatic PDAC^35^, we first confirmed that *HMGA2* is highly expressed in the basal subtype (**Fig. 1A**), and that *HMGA2*^+^ tumor spots associate with the presence of *FAP*^+^ CAFs (**Fig. 1B-C**) in the immediate neighborhood of primary tumors. Furthermore, *HMGA2*-positive tumor spots in metastatic liver and lung lesions similarly associated with increased expression of *FAP* compared to HMGA2 negative spots (**Fig. S1A**). To determine if tumor-derived factors were responsible for this fibroblast phenotype, we collected conditioned media from HMGA2^+^ basal (KP4, Panc3.27, BxPC3, and DanG) and HMGA2^-^ classical (SUIT2, HPAFII, SW1990, and Capan2) cell lines and applied it to pancreatic stellate cells (PSCs). HMGA2^+^ cell line conditioned media led to upregulation of *FAP* (**Fig. 1D**). In our genetically engineered mouse model (GEMM) of basal PDAC^17^, we found an increase in FAP^+^ fibroblasts in the TME of basal HMGA2^+^ mice with doxycycline-inducible LIN28B expression (*Kras^LSL-G12D/+^;LIN28B^LSL-Dox^;Ptf1a^Cre/+^, rtTA^+/-^* (K28C)) at disease endpoint compared to HMGA2^-^ littermates (*Kras^LSL-G12D/+^; Ptf1a^Cre/+^, rtTA^+/-^* (KC)) (**Fig. 1E, Fig. S1B**). Conditioned media from K28C-derived organoids was also sufficient to drive polarization towards *Fap*^+^ fibroblasts from naïve wild type pancreatic fibroblasts isolated from syngeneic mice (**Fig. 1F, Fig. S1C**), indicating tumor cell-intrinsic factors drive *Fap*^+^ fibroblast skewing downstream of HMGA2.

**Figure 1.**
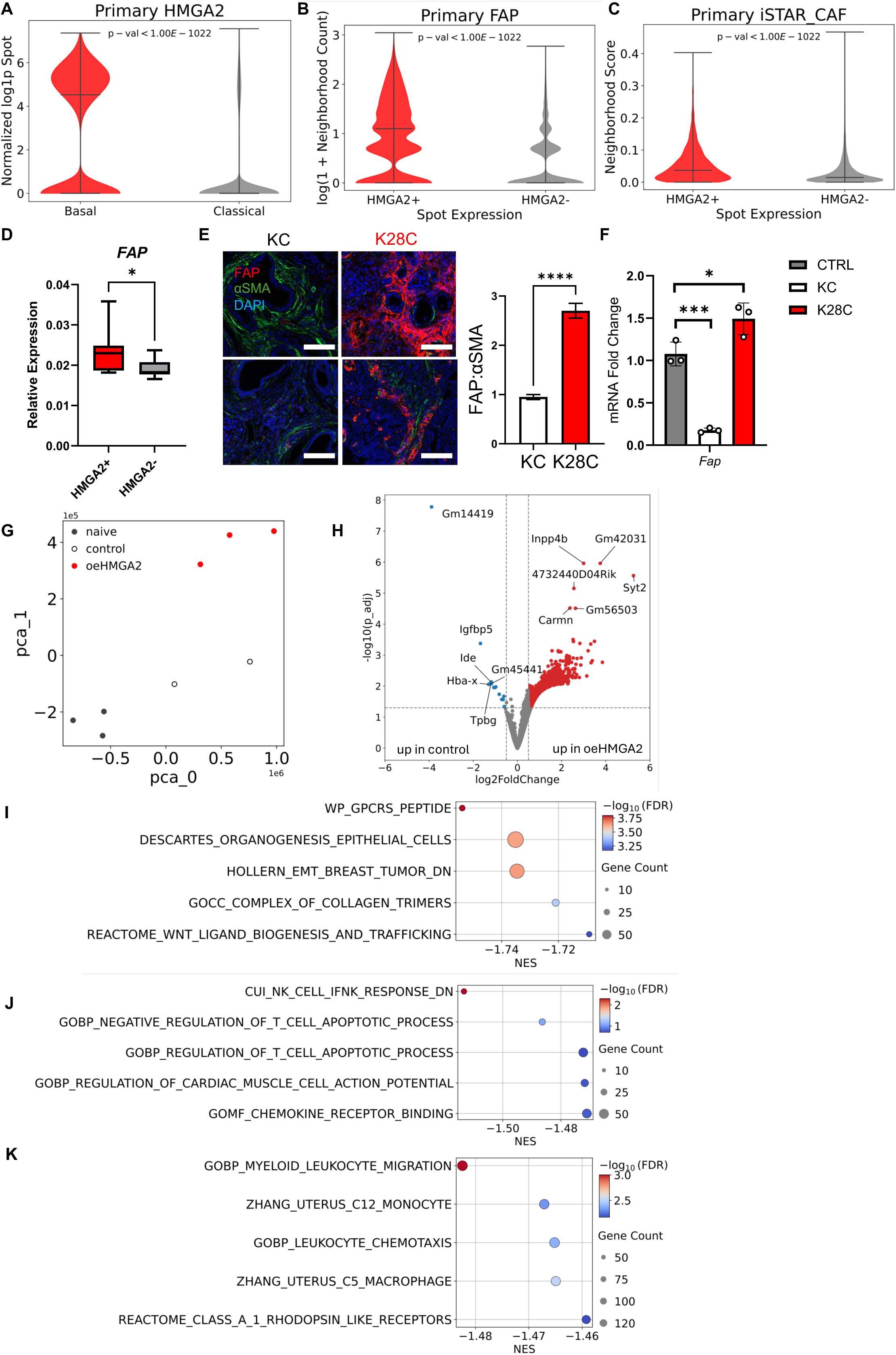
HMGA2^+^ basal PDAC tumors associate with altered fibroblast phenotypes. A) HMGA2 expression is higher in basal-subtyped tumor spots compared to classical-subtyped spots in 10x Visium spatial transcriptomics data. B) *FAP*^+^ areas are associated with *HMGA2*^+^ tumor spots in primary tumors from 10x Visium spatial transcriptomics data. C) *HMGA2*^+^ tumor spots have higher neighborhood cancer-associated fibroblast (CAF) signatures than *HMGA2*^-^ areas. D) Application of basal- or classical-conditioned media from 2D PDAC cell lines applied to naïve pancreatic stellate cells (PSCs) results in higher *FAP* expression in basal media-treated cells. n=4 cell lines per group (basal: KP4, Panc3.27, DanG, BxPC3; classical: SUIT2, HPAFII, SW1990, Capan2). E) Representative multiplex immunofluorescence (IF) on KC and K28C tumors at disease endpoint and quantification. Red = FAP, green = αSMA, blue = DAPI. Scale bar = 100 µm. F) Conditioned media from basal K28C tumor-derived organoids leads to higher *Fap* in naïve murine pancreatic fibroblasts compared to classical KC-derived organoids. G) PCA analysis of RNAseq on naïve pancreatic fibroblasts isolated from C57BL6/J mice treated with control media (grey), KPCY vector control media (white) or KPCY oeHMGA2 media (red). H) Volcano plot of upregulated genes in 2838 oeHMGA2 media-treated fibroblasts compared to KPCY vector control-treated fibroblasts. I) Five most downregulated GSEA signatures in KPCY vector control-treated fibroblasts. J) Five most downregulated GSEA signatures in KPCY oeHMGA2-treated fibroblasts. K) Five most downregulated GSEA signatures in K28C-treated fibroblasts. Data are graphed as mean ± SD, n ≥ 3 biological replicates unless otherwise indicated.

To determine the exact fibroblast phenotype driven by tumor cell *HMGA2* expression, we performed RNAseq on naïve murine pancreatic fibroblasts treated with conditioned media from the well-characterized 2838c3 KPCY (*Kras^LSL-G12D/+^;Trp53^LSL-R172H^/+;Pdx1-Cre;Rosa26^YFP/YFP^*) cell line developed by Li et al.^39^ engineered to overexpress *Hmga2* (**Fig. S1D**). Conditioned media from *Hmga2* expressing cells (oeHMGA2 CM) was associated with stronger variation away from naïve fibroblasts than control media-treated fibroblasts (**Fig. 1G**). Many more transcripts were upregulated in oeHMGA2 CM-treated fibroblasts than vector control-treated fibroblasts (**Fig. 1H**). When comparing both conditions back to baseline (**Fig. S1E-F**), control conditioned media led to downregulation of GSEA signatures for collagen genes and the Wnt signaling pathway (**Fig. 1I**), while oeHMGA2 CM led to downregulation of immune-related GSEA signatures (**Fig. 1J**). We performed parallel analyses on naïve fibroblasts treated with K28C conditioned media (**Fig. S1G**). Consistent with our oeHMGA2 findings, K28C media treatment downregulated immune-related pathways relative to naïve fibroblasts (**Fig. 1K**). Thus, HMGA2 expression *in vivo* associates with higher FAP^+^ CAF prevalence, and HMGA2^+^- conditioned media supports the development of an altered fibroblast phenotype *in vitro*.

### Tumoral HMGA2 expression increases IL6 and LRRC15 expression from adjacent fibroblasts and correlates with immunosuppression

Many subtypes of immunomodulatory CAFs have been described in PDAC, including inflammatory CAFs^25,26^, FAP^+^ CAFs^40^, CXCL12^+^ CAFs^41^, LRRC15^+^ CAFs^32^, and CXCL10^+^ CAFs^36^. Our data implicate a potential immunosuppressive role of HMGA2^+^ on CAFs in basal PDAC. We found that conditioned media from K28C GEMM-derived organoids and cell lines were sufficient to significantly increase *Il6* expression in treated naïve fibroblasts (**Fig. 2A-B**). Downregulating HMGA2 levels by removing doxycycline from tumor cells also led to lower *Il6* expression in treated fibroblasts (**Fig. 2B**). Conditioned media from basal PDAC cell lines similarly led to increased *IL6* and *LRRC15* expression compared to classical conditioned media in treated PSCs (**Fig. 2C**). shRNA-mediated knockdown of *HMGA2* in a panel of basal PDAC cell lines was sufficient to decrease *IL6* and *LRRC15* expression, but not *ACTA2* or *FAP*, in conditioned media-treated PSCs (**Fig. 2D, Fig. S2A**). In our basal PDAC GEMM, KC and K28C tumors had equivalent PDPN^+^ fibroblasts, but K28C tumors had increased IL6^+^ CAFs (**Fig. 2E, Fig. S2B**). As both IL6^+^ and LRRC15^+^ CAF populations have been described to trigger immune suppression in PDAC and other cancers^27,32,42^, we performed IHC and multiplex flow cytometry on K28C and KC tumors. Large-scale lymphoid alterations were detected in HMGA2^+^ tumors, including poor CD8+ T cell infiltration detected by IHC (**Fig. 2F, Fig. S2C**), and decreased effector CD44^+^ CD62L^-^ CD8^+^ T cells and decreased effector CD44^+^ CD62L^-^ CD4^+^ T cells detected by flow cytometry (**Fig. 2G**). In contrast, Ly6G^+^, Ly6C^+^, and F4/80^+^ populations were unaffected by high expression of *Hmga2* (**Fig. S2D-F**). These findings were recapitulated in KPCY allografts, where expression of *Hmga2* was significantly higher in T cell-low clones compared to T cell-high clones (**Fig. 2H**)^39^. Additionally, orthotopic injection of T cell-low lines induced a higher abundance of LRCC15^+^ fibroblasts than T cell-high cell lines (**Fig. S2G**). Analysis of spatial transcriptomics datasets also showed HMGA2^+^ tumor cell-enriched spots associated with fewer CD8^+^ T cells (**Fig. S2H**). Further, analysis of publicly available scRNAseq data from human PDAC tumors^36^ revealed fewer CD8^+^ T cells and more Tregs in HMGA2^+^ tumors compared to HMGA2^-^ tumors (**Fig. 2I**).

**Figure 2.**
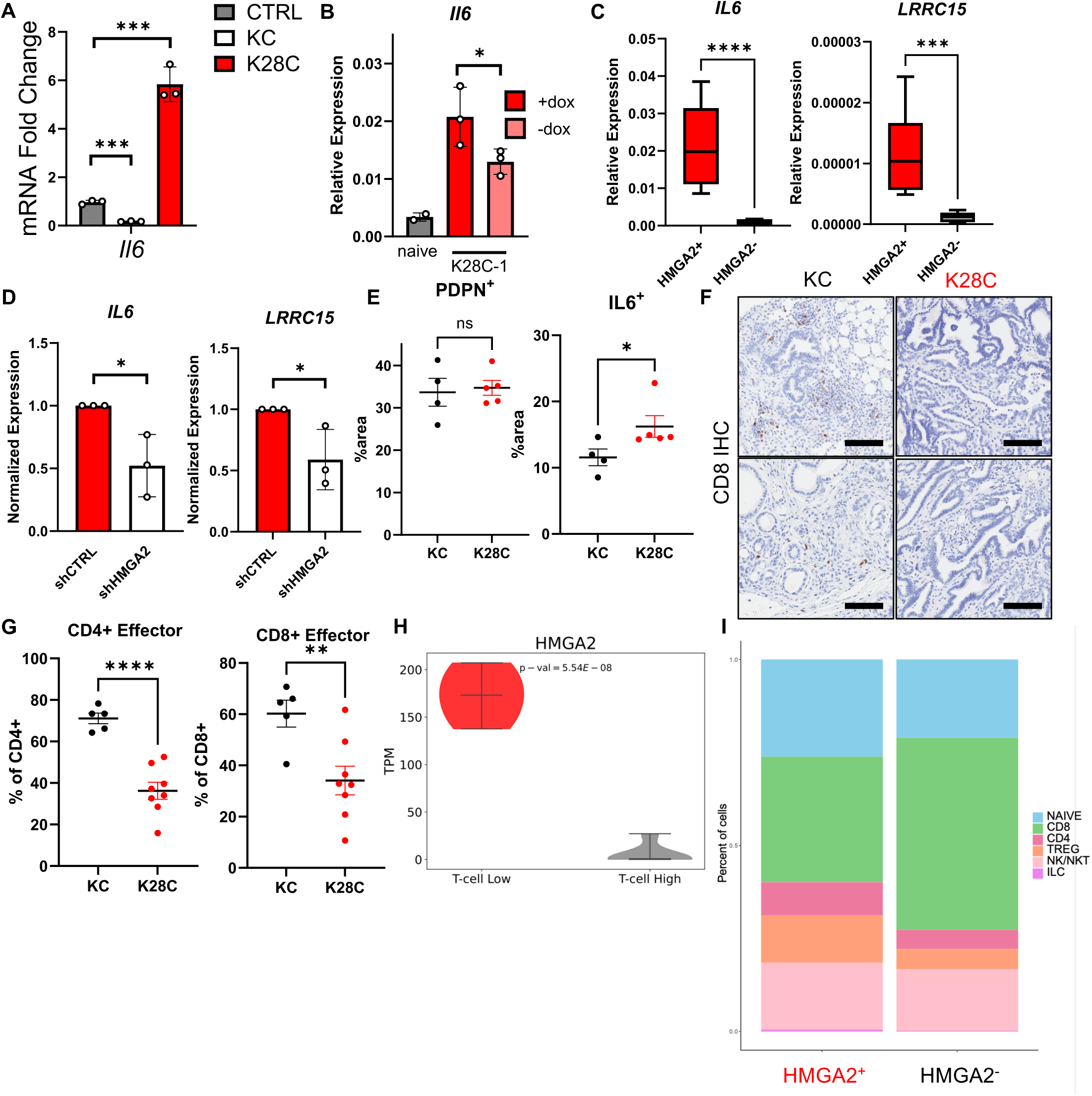
HMGA2 expression in tumor cells leads to upregulation of IL6 in adjacent fibroblasts and functional immunosuppression. A) Application of conditioned media from K28C-derived organoids to naïve murine pancreatic fibroblasts results in upregulation of *Il6*. B) Removal of doxycycline (dox) treatment from K28C-derived 2D cell lines leads to decreased *Il6* expression in treated fibroblasts compared to dox on control. C) Application of basal- or classical-conditioned media from 2D PDAC cell lines applied to naïve pancreatic stellate cells (PSCs) results in higher *IL6* and *LRRC15* in basal media-treated cells. n = 4 cell lines per group (basal: KP4, Panc3.27, DanG, BxPC3; classical: SUIT2, HPAFII, SW1990, Capan2). D) shRNA-mediated knockdown of *HMGA2* in three basal cell lines (KP4, Panc1, Panc3.27) leads to downregulation of *IL6* and *LRRC15* in treated human PSCs compared to control. E) IF on K28C and KC primary tumors reveals higher fibroblast IL6 in K28C animals. n = 5 per group. F) Immunohistochemistry (IHC) on K28C and KC tumors shows decreased CD8+ T cell infiltrate into K28C tumors compared to KC. Scale bar = 100 µm. G) Flow cytometry on endpoint K28C and KC tumors shows a relative paucity of CD44^+^ CD62L^-^ CD4^+^ and CD44^+^ CD62L^-^ CD8^+^ effector T cells in K28C tumors. n(KC) = 5; n(K28C) = 8. H) RNAseq on previously published KPCY cell line clones shows enrichment for HMGA2 in T cell-low clones. I) scRNAseq atlas analysis reveals greater proportion of Tregs and smaller proportion of CD8^+^ T cells in HMGA2^+^ tumors, with all cell types graphed significantly altered between HMGA2^+^ and HMGA2^-^ groups. Data are graphed as mean ± SD, n ≥ 3 biological replicates unless otherwise indicated.

### HMGA2 controls expression of IL1A

We then conducted a candidate gene analysis for enriched cytokines and chemokines in HMGA2^+^ basal cells (KEGG pathway hsa04060) in three independent PDAC transcriptomic datasets: (1) bulk RNAseq from 79 patient samples at the University of Pennsylvania; (2) laser capture microdissected RNAseq from the COMPASS trial; and (3) single cell RNAseq^36^ (**Fig. 3A-B, Fig. S3A-C**). Only two genes were consistently and significantly upregulated in HMGA2^+^ tumors: *IL1A* and *IL20RB*. TGFβ family members *TGFB1* and *TGFB2* were not consistently enriched in HMGA2^+^ tumors (**Fig. 3A, Fig. S3A-C**). Since the TGFβ and IL1 families of cytokines are both heavily implicated in controlling fibroblast phenotypes in PDAC^25,26,32^, we next tested whether HMGA2 controls the levels of these cytokines. In subtyped 2D PDAC cell cultures, HMGA2 expression was associated with significantly higher levels of *IL1A*, but not *IL1B*, *TGFB1*, or *TGFB2* (**Fig. 3C**). Furthermore, knockdown of *HMGA2* in basal lines led to significant reduction in *IL1A* transcription, with no consistent effects on *IL1B*, *TGFB1,* or *TGFB2* (**Fig. 3D, Fig. S3D**). Conversely, doxycycline-induced overexpression of *HMGA2* in classical cell lines increased *IL1A* transcription without altering *IL1B* or *TGFB1* levels (**Fig. 3E, Fig. S3E**). *HMGA2* knockdown also decreased IL1α in whole cell lysates (**Fig. 3F**) and cell culture supernatant (**Fig. 3G**), confirming that modulation of HMGA2 is sufficient to alter intracellular and extracellular IL1α levels. Withdrawal of doxycycline from K28C cell lines led to reduction of *Hmga2* and *Il1a* (**Fig. 3H, Fig. S3F**), as well as decreased secretion of IL1α into the media (**Fig. 3I**). In the KPCY mouse model, overexpression of *Hmga2* increased *Il1a* transcription and IL1α levels in media (**Fig. 3J-K, Fig. S3G**), while doxycycline inducible shRNA-mediated knockdown of *Hmga2* decreased *Il1a* transcription and IL1α levels in media (**Fig. 3L-M, Fig. S3H**). In both systems, *Tgfb1* and *Cxcl1*^39^ expression was not significantly altered (**Fig. S3G-H**).

**Figure 3.**
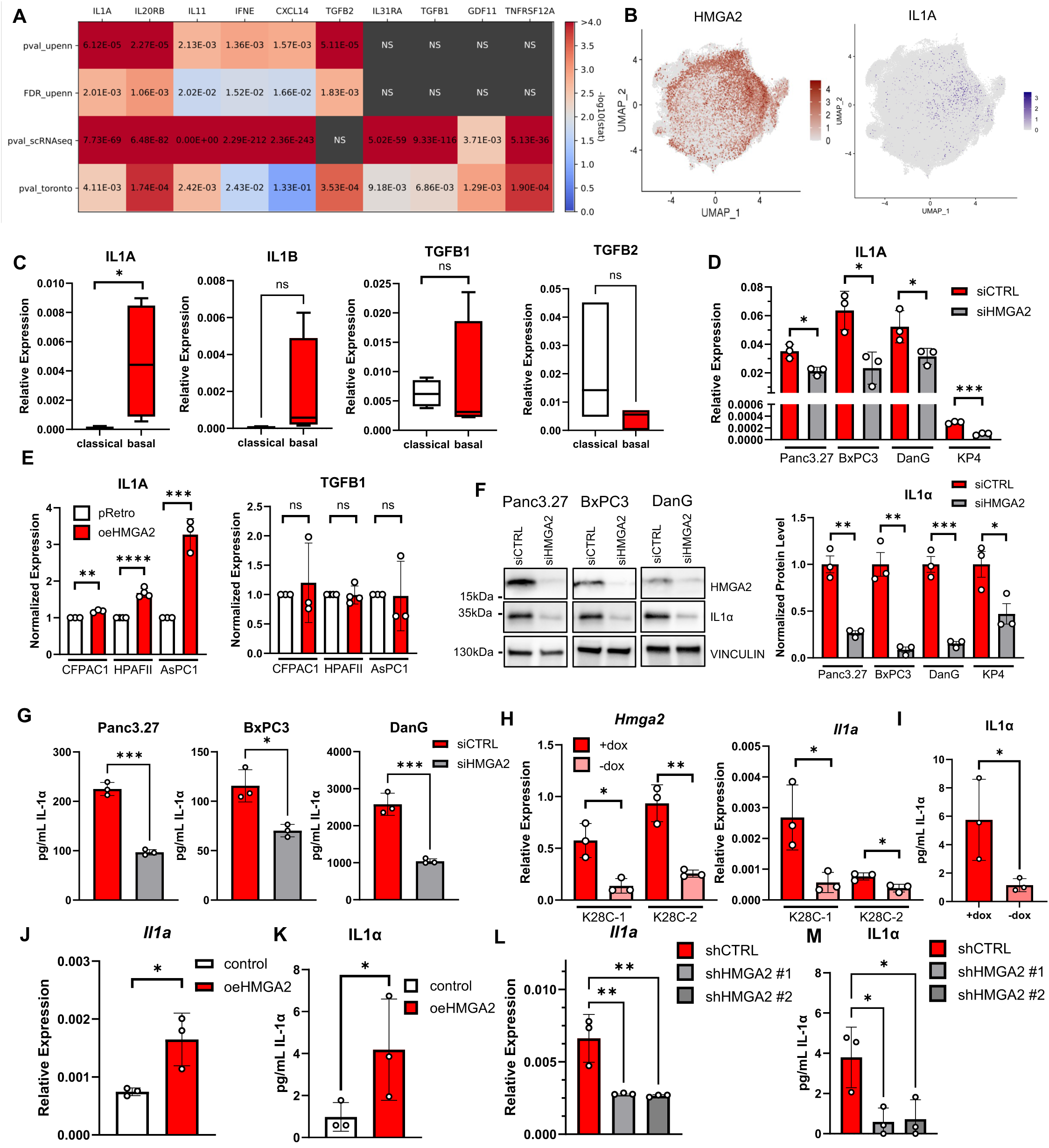
HMGA2 levels control IL1A expression in basal PDAC. A) Heatmap of significance of association between HMGA2^+^ tumor samples and indicated cytokine genes from three individual datasets. UPenn = bulk RNAseq (n=79), Toronto = COMPASS trial. B) scRNAseq UMAP of ductal cell object with expression of *HMGA2* and *IL1A* expression overlaid. C) Baseline expression of *IL1A*, but not *IL1B*, *TGFB1*, or *TGFB2* is higher in basal cells than in classical cells. n = 4 cell lines per group (basal: KP4, Panc3.27, DanG, BxPC3; classical: SUIT2, HPAFII, SW1990, Capan2). D) siRNA-mediated knockdown of *HMGA2* in four basal cell lines leads to reduced transcription of *IL1A*. E) Overexpression of HMGA2 in three classical cell lines increases *IL1A* but not *TGFB1* transcription. F) Western blot of whole cell lysates with and without knockdown of *HMGA2* shows that loss of HMGA2 leads to decreased intracellular IL1α. Quantified on right. G) Enzyme-linked immunosorbent assay (ELISA) of conditioned media from siCTRL and siHMGA2 cell lines shows HMGA2 loss decreases extracellular IL1α. H) Dox withdrawal from K28C cell lines leads to decreased *Hmga2* and *Il1a* expression. I) ELISA on K28C-conditioned media shows decreased IL1α with dox withdrawal. J) Overexpression of *Hmga2* in a T cell-high KPCY cell line leads to increased *Il1a* expression. K) ELISA shows increased IL1α in cell culture media from KPCY cells with *Hmga2* overexpression. L) Dox-inducible shRNA mediated knockdown of *Hmga2* in a T cell-low KPCY cell line leads to decreased *Il1a* expression. M) ELISA shows decreased IL1α in cell culture media from KPCY cells with *Hmga2* knockdown. Data are graphed as mean ± SD, n ≥ 3 biological replicates unless otherwise indicated.

### HMGA2 binds the *IL1A* promoter and displaces linker histone H1

To determine if HMGA2 directly influences *IL1A* transcription, we performed chromatin immunoprecipitation qPCR (ChIP-qPCR) at the *IL1A* promoter in four basal cell lines. HMGA2 was detected binding ∼412 base pairs before the *IL1A* transcription start site (TSS) (**Fig. 4A, Fig. S4A**). siRNA-mediated knockdown of *HMGA2* abrogated binding and led to decreased trimethylation of histone 3 lysine 4 (H3K4me3) at the same locus (**Fig. 4A**). HMGA2 has previously been reported to compete with the linker histone H1 for DNA binding, leading to either transcriptional repression or activation^17,43,44^. We found that knockdown of HMGA2 increased H1 occupancy at the *IL1A* promoter, specifically the histone variants H1.1 and H1.5 (**Fig. 4B**). The H1 linker histone is responsible for chromatin compaction on the core nucleosome, and HMGA2 has been previously implicated in displacing the repressive H1.5 variant^45^. These data provide evidence that HMGA2 remodels the chromatin landscape directly surrounding the *IL1A* TSS.

**Figure 4.**
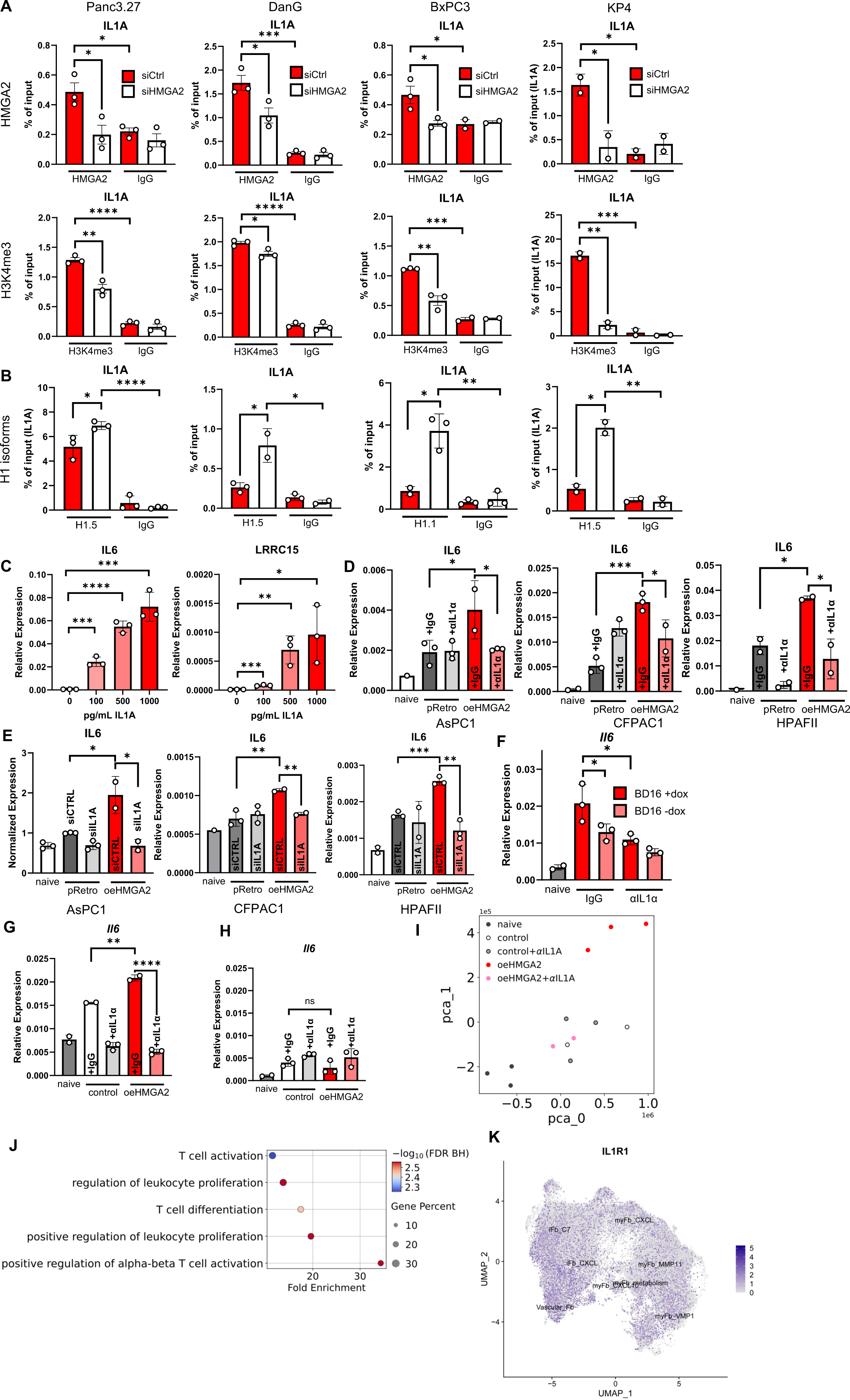
HMGA2 controls chromatin landscape at the IL1A promoter and is sufficient to drive fibroblast polarization. A) ChIP-qPCR in four basal cell lines shows HMGA2 binding at the *IL1A* promoter, with high H3K4me3. siHMGA2 abrogates HMGA2 binding and results in decreased H3K4me3. B) HMGA2 knockdown leads to increased H1.1/H1.5 at the IL1A promoter. Recruited H1 isotype is cell-line specific. C) Treatment of naïve PSCs with concentrations of IL1α equivalent to amounts released by basal cells is sufficient to increase *IL6* and *LRRC15* transcription in a dose-dependent manner. D) Conditioned media from three HMGA2-overexpressing classical cell lines is sufficient to drive increased *IL6* in treated PSCs compared to empty vector control. Neutralizing IL1α reverses this effect. E) siRNA-mediated knockdown of *IL1A* in tumor cells is sufficient to reverse HMGA2-driven fibroblast skewing when conditioned media is applied to naïve PSCs. F) Treatment of K28C conditioned media with IL1α neutralizing antibody prevents HMGA2-driven fibroblast skewing. G) Conditioned media from an HMGA2-overexpressing KPCY cell line is sufficient to drive increased *Il6* in treated naïve murine pancreatic fibroblasts compared to empty vector control. Neutralizing IL1α reverses this effect. H) Treating *Il1r1* knockout fibroblasts with oeHMGA2 conditioned media does not alter fibroblast polarization. I) PCA plot showing fibroblasts treated with αIL1α cluster with vector control-treated fibroblasts. J) Top five GO terms enriched in oeHMGA2-conditioned fibroblasts when treated with anti-IL1α. K) scRNAseq atlas analysis of cancer-associated fibroblast UMAP object with overlaid IL1R1 expression. Data are graphed as mean ± SD, n ≥ 2 biological replicates unless otherwise indicated.

### HMGA2-driven IL1A is sufficient to drive an immunosuppressive fibroblast phenotype

We then tested whether activation of IL1α/IL1R1 by HMGA2 was sufficient to drive fibroblast skewing. We found that the levels of IL1α produced by basal cell lines were sufficient to upregulate *IL6*, *LRRC15, FAP,* and *TGFB2* transcription in treated PSCs (**Fig. 4C, Fig. S4B**). Meanwhile, *ACTA2* levels remained consistent (**Fig. S4B**). When we applied conditioned media from classical cell lines overexpressing *HMGA2* to naïve PSCs, we found that *HMGA2* overexpression was sufficient to drive a significant increase in fibroblast *IL6* transcription and was completely reversible by treatment with an anti-IL1α neutralizing antibody (**Fig. 4D, Fig. S4C**). siRNA-mediated knockdown of *IL1A* in PDAC cell lines was also sufficient to reverse fibroblast *IL6* polarization by conditioned media (**Fig. 4E, Fig. S4D**). Doxycycline withdrawal from K28C mouse lines was sufficient to decrease *Il6* when conditioned media was applied to naïve pancreatic fibroblasts, as was treatment with anti-IL1α (**Fig. 4F, Fig. S4E**). Conditioned media from the KPCY T cell-low cell line 2838c3 with overexpression of *Hmga2* also increased *Il6* transcription when applied to naïve pancreatic fibroblasts, and this skewing was reversed when conditioned media was treated with anti-IL1α antibodies (**Fig. 4G, Fig. S4F**) or when conditioned media was applied to naïve pancreatic fibroblasts isolated from mice lacking the IL1 receptor (*Il1r1*^-/-^) (**Fig 4H, Fig. S4G**). RNAseq showed that IL1α blockade normalized oeHMGA2-treated fibroblasts back to vector control treated fibroblasts (**Fig. 4I)** and lead to massive upregulation of gene pathways involved in leukocyte activation and differentiation (**Fig. 4J, Fig. S4H).** We also confirmed that *IL1R1* is ubiquitously expressed across CAF subtypes in scRNAseq in patient tumors, demonstrating that this pathway can be activated regardless of CAF subtype (**Fig. 4K**)^36^.

### HMGA2 overexpression leads to immunosuppression *in vivo*

We were then motivated to determine if HMGA2-driven fibroblast skewing was sufficient to cause functional alterations *in vivo* in an immunocompetent setting. Orthotopic injection of KPCY cells with *Hmga2* overexpression into syngeneic wild-type mice led to increased tumor mass and increased liver and lung metastases compared to empty vector control (**Fig. 5A, Fig. S5A**). Fibroblast alterations were apparent at the tumor margin (**Fig. 5A**). High HMGA2 expression was maintained *in vivo* and observed in lung metastases (**Fig. 5B, Fig. S5B**). Flow cytometry revealed decreased intratumoral CD3^+^ T cells, CD8^+^ cytotoxic T cells, NK cells, and Type-1 conventional dendritic cells (cDC1) in oeHMGA2 tumors (**Fig. 5C, Fig. S5C-D, Table S1**). Total CD4^+^ T cells and CD4^+^ FoxP3^+^ CD25^+^ regulatory T cells (Tregs) were increased in oeHMGA2 tumors (**Fig. 5C, Fig. S5C**). Total CD45^+^ infiltrate, CD11b^+^ MHCII^-^ cells, neutrophils, Type-2 conventional dendritic cells (cDC2), monocytes, and macrophages were not significantly altered by oeHMGA2 (**Fig. S5C**). Multiplex immunofluorescence for FAP/αSMA revealed a slight increase in αSMA^+^ fibroblasts and a marked increase in FAP^+^ fibroblasts in oeHMGA2 tumors (**Fig. 5D**). Furthermore, overall PDPN^+^ fibroblast content, IL6^+^ fibroblast content, and LRRC15^+^ fibroblast content were higher with HMGA2 overexpression (**Fig. 5E-F**). We further confirmed that these immunologic changes were due to IL1α/IL1R1 signaling by implanting orthotopic tumors into *Il1r1*^-/-^ mice, where we observed similar growth kinetics but no association with lung metastasis or tumor-edge fibrosis on histology (**Fig. S5E-F**). At endpoint, intratumoral CD3^+^ T cells remained lower in oeHMGA2 tumors, indicating that other signaling pathways besides IL1α/IL1R1 play a role in stimulating total T cell infiltrate (**Fig. 5G**). However, CD8^+^ cytotoxic T cells, NK cells, CD4^+^ T cells, and Tregs were no longer significantly different between control and oeHMGA2 tumors in *Il1r1*^-/-^ mice, demonstrating that signaling via the IL1α/IL1R1 axis is required for the lymphocyte-driven immunosuppression found in HMGA2^+^ tumors (**Fig. 5G**). Meanwhile, the proportion of CD11b^+^ MHC-II^-^ cells increased in *Il1r1*^-/-^ animals compared to wild-type animals, indicating that other cytokine signaling programs drive a myeloid-predominant phenotype in the absence of IL1 signaling (**Fig. S5G**). In the tumor stroma, FAP^+^, IL6^+^, and LRRC15^+^ CAFs were not induced with full-body *Il1r1* knockout, while αSMA^+^ fibroblasts remained higher in oeHMGA2 tumors (**Fig. 5H-I, Fig. S5H**). These results implicate HMGA2-driven IL1α/IL1R1 signaling in driving immunosuppressive fibroblast polarization and in controlling immune cell infiltrate in the basal PDAC TME.

**Figure 5.**
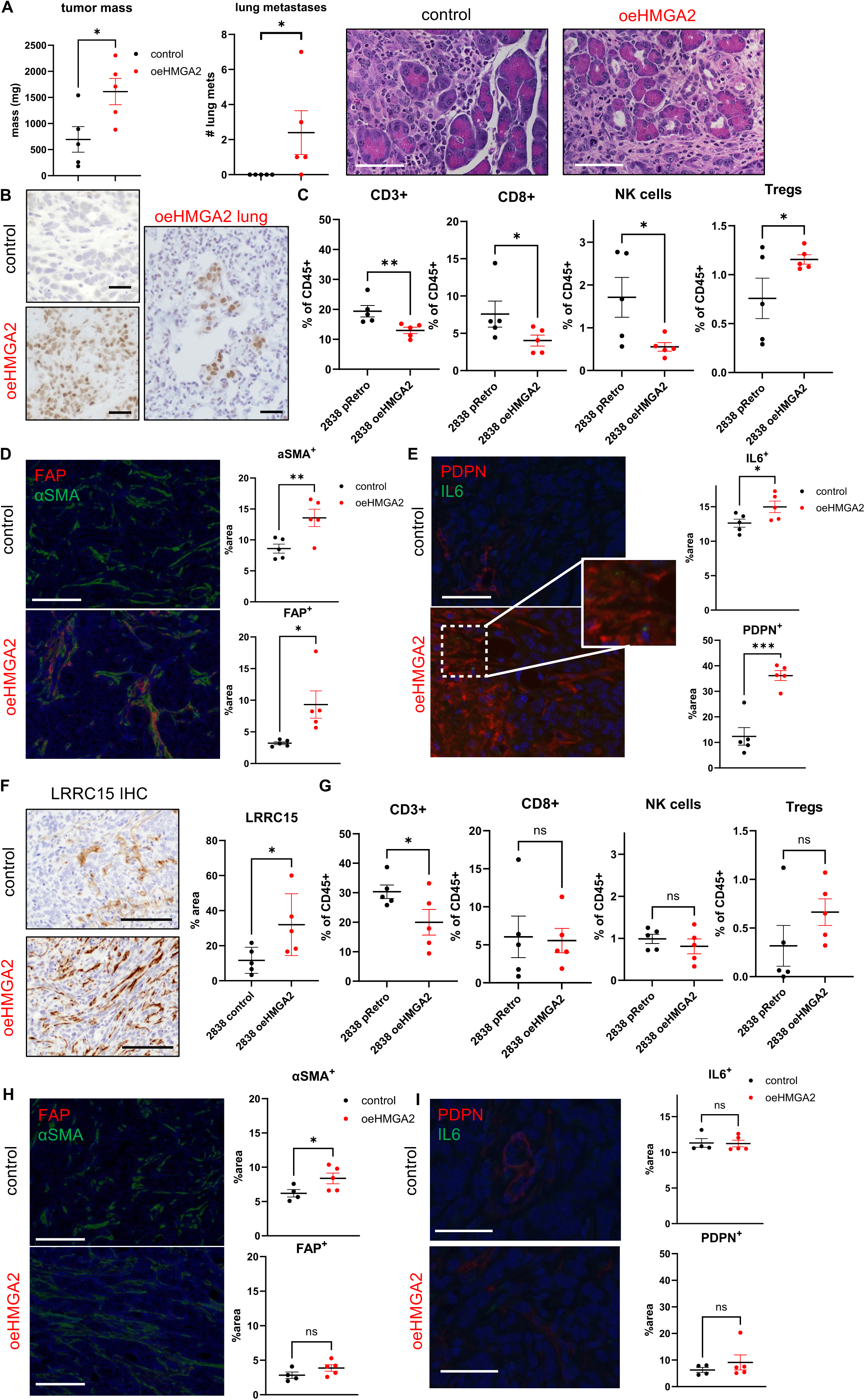
HMGA2 drives fibroblast polarization and immunosuppression *in vivo* via IL1α. A) Orthotopic injection into syngeneic pancreas of KPCY oeHMGA2 cells increases final tumor mass, lung metastases, and leading-edge fibrosis compared to empty vector control. Scale bar = 50 µm. B) HMGA2 IHC on vector control and oeHMGA2 tumors and associated oeHMGA2 lung metastasis. Scale bar = 25 um. C) Flow cytometry on endpoint tumors reveals decreased CD3^+^ T cells, CD8^+^ cytotoxic T cells, and NK cells with tumoral HMGA2 expression. CD25^+^ FoxP3^+^ T regs increase in oeHMGA2 tumors. D) IF shows increased αSMA and FAP in cancer-associated fibroblasts with oeHMGA2. Scale bar = 50 µm. E) Total fibroblast content (PDPN^+^) and IL6^+^ fibroblasts increase with oeHMGA2. Scale bar = 50 µm. F) oeHMGA2 tumors have higher LRRC15^+^ fibroblast content than control tumors. Scale bar = 50 um. G) Flow cytometry on endpoint tumors in IL1R1 knockout shows decreased CD3^+^ T cells in oeHMGA2 tumors. CD8^+^ cytotoxic T cell, NK cell, and Treg cell prevalence is no longer differential based on HMGA2 expression. H) IF reveals aSMA levels in CAFs increases with oeHMGA2 but FAP remains flat when host cells lack IL1R1. Scale bar = 100 µm. I) Total fibroblasts and IL6^+^ fibroblasts levels are similar in empty vector and oeHMGA2 tumors when host cells lack IL1R1. Scale bar = 50 µm. Data are graphed as mean ± SEM, n = 5 per group.

### IL1R1 on both immune and stromal cells is necessary for HMGA2-induced immunosuppression

Our data demonstrates that the IL1α/IL1R1 axis is necessary for decreased CD8^+^ T cell and NK cell infiltrate, as well as increased Tregs, into HMGA2^+^ PDAC tumors. However, it remained unclear if IL1α released by tumor cells was signaling directly to these immune cells or if IL1R1 activation on fibroblasts and downstream upregulation of IL6 and other cytokines led to the TME alterations observed. We therefore generated bone marrow chimeras by reconstituting *Il1r1*^-/-^ recipients with WT bone marrow and WT recipients with *Il1r1*^-/-^ bone marrow, then orthotopically implanted these mice with KPCY cells with and without overexpression of *Hmga2* (**Fig. 6A, Fig. S6A-B**). At endpoint, all conditions led to no difference in tumor mass or lung metastases between control and oeHMGA2 tumors (**Fig. 6B, Fig. S6C**). Interestingly, neither CD8^+^ T cell nor NK cell differences were maintained between control and oeHMGA2 tumors when IL1R1 was lost on either immune or non-immune cells (**Fig. S6D-E**). However, Treg differences based on tumor subtype were maintained in mice with an *Il1r1* knockout immune system (**Fig. 6C**). Therefore, Treg differences between HMGA2^+^ and HMGA2^-^ tumors are not reliant on IL1 signaling directly to immune cells, even though IL1R1^+^ Tregs have been previously reported to be pro-tumorigenic^46^. Instead, an intermediary non-hematopoietic cell induces Treg recruitment in HMGA2^+^ tumors. IL1R1 signaling is required in both hematopoietic and non-hematopoietic cells to observe differences in CD8^+^ T cell and NK cell infiltrate depending on tumor subtype.

**Figure 6.**
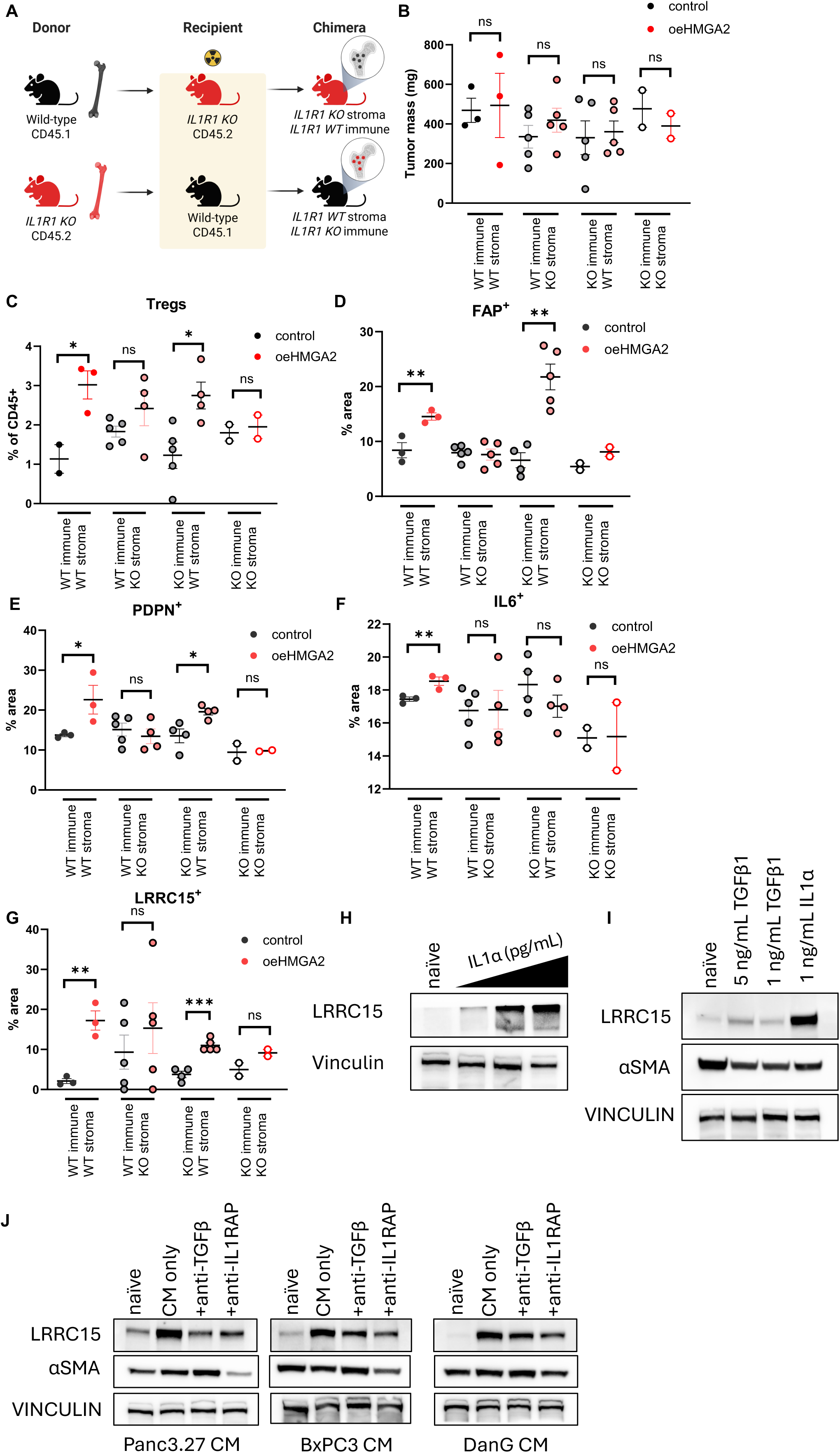
IL1A does not directly signal immune cells to result in immunosuppression. A) Schema for syngeneic bone marrow chimera, leading to selective knockout of IL1R1 on hematopoietic or non-hematopoietic cells. B) Chimera mice tumors do not grow as large as non-irradiated studies. Tumor masses are not different between vector control and oeHMGA2 tumors when IL1R1 is missing on either immune or stromal cells. C) Knockout of IL1R1 on immune cells allows for differential Treg recruitment between control and oeHMGA2 tumors, while WT immune cells with stromal IL1R1 KO leads to abrogation of this effect. D) FAP^+^ fibroblasts are significantly affected by tumoral HMGA2 expression only when fibroblasts have WT IL1R1. E) PDPN^+^ fibroblasts are significantly affected by tumoral HMGA2 expression only when fibroblasts have WT IL1R1. F) KO of IL1R1 on either stromal or immune cells prevents IL6 upregulation in CAFs by tumoral HMGA2 expression. G) LRRC15^+^ CAFs are increased in oeHMGA2 tumors when fibroblasts have WT IL1R1. H) Treatment of naïve PSCs with increasing doses of recombinant IL1α leads to increased LRRC15 protein levels. I) IL1α induces LRRC15 expression in treated PSCs more than recombinant TGFβ treatment. J) Neutralization of IL1RAP or TGFβ in basal conditioned media leads to reduced LRRC15 levels in PSCs. Data are graphed as mean ± SEM, n ≥ 2 per group.

In the non-hematopoietic compartment, *Il1r1* knockout was sufficient to abrogate HMGA2-mediated FAP^+^ (**Fig. 6D**) and PDPN^+^ (**Fig. 6E**) fibroblast increases in the TME. Differences in fibroblast IL6 levels, however, were dependent on IL1R1 presence on both the hematopoietic and non-hematopoietic compartments (**Fig. 6F**), and aSMA^+^ fibroblasts trended higher when IL1R1 was present on only immune cells (**Fig. S6F**). Finally, IL1R1 was required on non-hematopoietic cells to see a significant difference in LRRC15^+^ fibroblast content between HMGA2^+^ and HMGA2^-^ tumors (**Fig. 6G**). Therefore, HMGA2^+^ tumor cell-driven IL1α signaling to fibroblasts contributes to upregulation of FAP, PDPN, and LRRC15 in neighboring fibroblasts, but IL1R1 signaling in immune cells plays a role in controlling αSMA and IL6 expression. Interestingly, LRRC15^+^ fibroblasts were enriched in control tumors with *Il1r1* KO immune systems (**Fig. 6G**), indicating that IL1R1 signaling on immune cells may suppress LRRC15^+^ fibroblasts in the immunocompetent setting. These data emphasize the importance of stromal and mesenchymal IL1R1 signaling in controlling the immunophenotype of basal PDAC tumors.

### IL1α directly induces LRRC15 expression in PSCs

LRRC15^+^ fibroblasts have been previously described in multiple human cancer types and are commonly found in PDAC^32^. Previous work by Krishnamurty et al. found that LRRC15^+^ CAFs support a fibroblast-rich TME, limit CD8^+^ T cell infiltration, and prevent responses to immune checkpoint blockade^32^. However, their work focuses on TGFβ signaling, which would not explain the difference in LRRC15^+^ CAFs we observe between HMGA2^+^ and HMGA2^-^ tumors. When we treated naïve human PSCs with recombinant IL1α, we found that doses as low as 100 pg/mL were sufficient to induce *LRRC15* and *TGFB2* transcription (**Fig. 4C**) and to increase LRRC15 protein levels (**Fig. 6H**). Furthermore, LRRC15 levels were higher when PSCs were treated with IL1α compared to TGFβ1 or TGFβ2 (**Fig. 6I, Fig. S6G-H**). When PSCs are treated with conditioned media from basal HMGA2^+^ cell lines, the baseline increases in LRRC15 and αSMA are abrogated by blockade of IL1 receptor accessory protein (IL1RAP), the necessary dimerization protein of IL1R1, equally or more efficiently than blockade of TGFβ1-3 (**Fig. 6J, Fig. S6I**). These data support the hypothesis that HMGA2-driven IL1α signaling plays a parallel and complementary role to TGFβ-driven upregulation of LRRC15 in naïve fibroblast precursors.

### Targeting IL1RAP in basal PDAC

The IL1 signaling axis has been extensively targeted both in autoimmune disease and cancer^47–49^. Direct antagonism of IL1R1 in PDAC with anakinra has been attempted with limited success (NCT02021422, NCT02550327). Nadunolimab, a fully humanized antibody targeting IL1RAP, has shown efficacy and tolerability in locally advanced/metastatic PDAC in combination with gemcitabine and nab-paclitaxel, particularly in patients with high tumoral IL1RAP expression at baseline^50,51^. High levels of IL1RAP also predict poorer overall survival in PDAC^52^. We found that IL1RAP is significantly higher in HMGA2^+^ basal PDAC by bulk RNAseq (**Fig. 7A**). *IL1RAP* was also highly expressed in the vicinity of HMGA2^+^ tumor spots across primary and metastatic lesions by spatial transcriptomics (**Fig. 7B**). Functionally, anti-IL1RAP treatment was sufficient to reverse *IL6* increases in oeHMGA2 conditioned media-treated human PSCs (**Fig. 7C, Fig. S7A**) and naïve murine fibroblasts (**Fig. 7D, Fig. S7B**). When basal-derived conditioned media was applied to PSCs, treatment with nadunolimab decreased levels of both *IL6* and *LRRC15* (**Fig. 7E**, **Fig. 6J, Fig. S6I**). RNAseq analysis of anti-IL1RAP-treated fibroblasts treated with oeHMGA2 conditioned media revealed that blocking IL1RAP once again restored fibroblasts back to a baseline gene expression signature (**Fig. 7F**) and led to upregulation of immune response-related GO pathways (**Fig. S7C**). *In vivo,* treatment of oeHMGA2 KPCY tumors with murine anti-IL1RAP was well tolerated and resulted in no difference in final tumor mass (**Fig. S7D**). However, the number of lung metastases from oeHMGA2 tumors was strongly decreased with anti-IL1RAP treatment (**Fig. 7G**). In the TME, anti-IL1RAP led to a decrease in FAP^+^ fibroblasts and an increase in αSMA^+^ fibroblasts, with limited effects on PDPN^+^ fibroblasts (**Fig. 7H, Fig. S7E-F**). LRRC15^+^ and IL6^+^ fibroblasts were significantly decreased in oeHMGA2 tumors with anti-IL1RAP (**Fig. 7I**). Treg presence in tumors was also decreased by anti-IL1RAP treatment, with no significant alterations in CD3^+^, CD4^+^, or CD8^+^ T cells or CD11b^+^ MHCII^-^ myeloid content (**Fig. 7J, Fig. S7G**). Therefore, treatment with anti-IL1RAP allows for targeting of the HMGA2/IL1α/fibroblast axis in basal PDAC, leading to decreased Tregs and LRRC15^+^ fibroblasts in the TME, which may contribute to decreased lung metastases.

**Figure 7.**
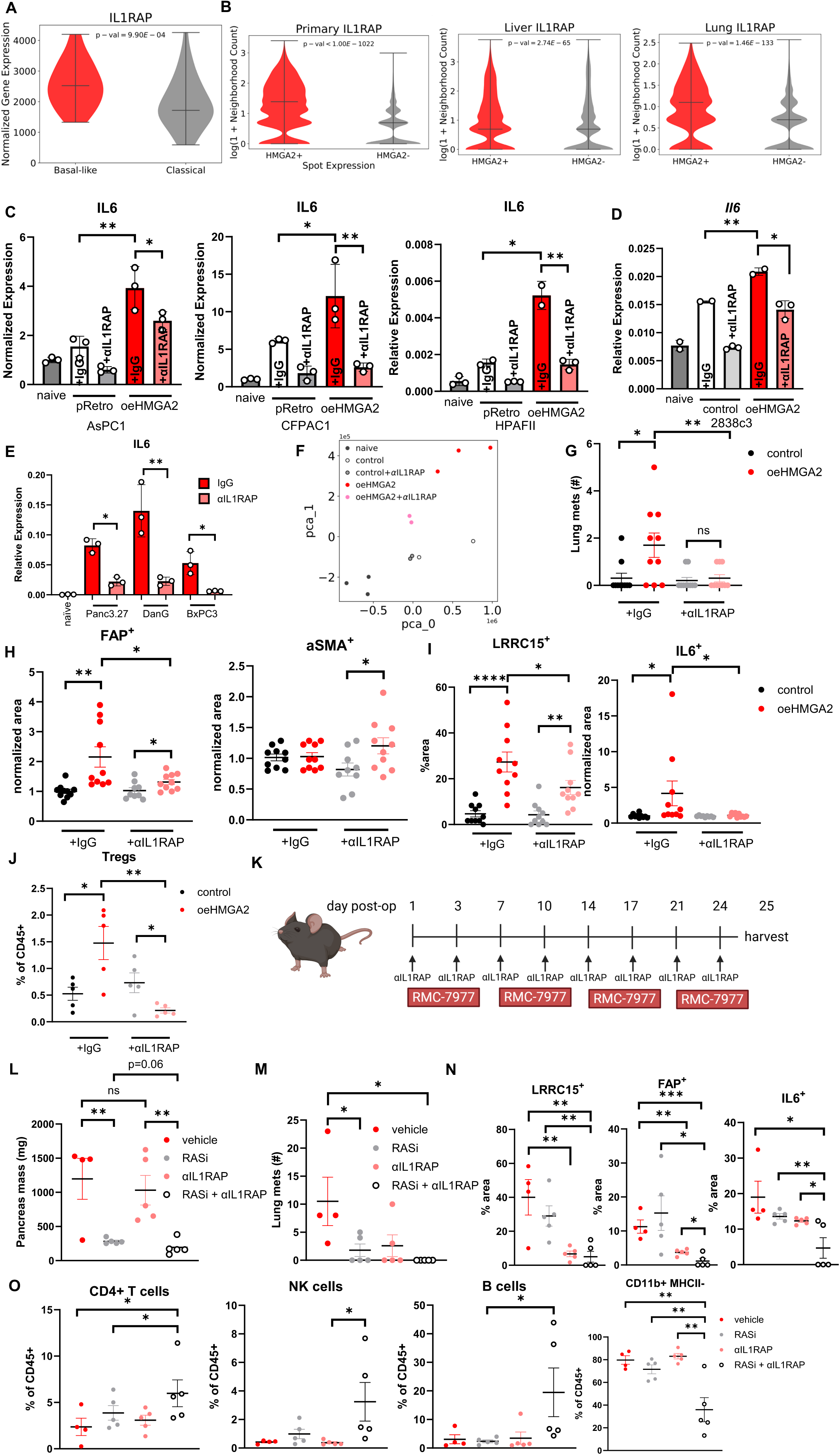
Blocking IL1RAP allows for normalization of the TME and increases efficacy of KRAS inhibition. A) Bulk RNAseq from 79 patients shows higher IL1RAP expression in basal PDAC tumors. B) Spatial transcriptomics studies confirm higher IL1RAP in the vicinity of HMGA2^+^ tumor spots in primary tumors, liver metastases, and lung metastases. C) Treatment of conditioned media with nadunolimab reverses *IL6* increases in treated PSCs seen with HMGA2 overexpression. D) Treatment of conditioned media from 2838c3 cell lines with mouse anti-IL1RAP decreases *Il6* in oeHMGA2-treated fibroblasts back to baseline levels. E) Pancreatic stellate cells upregulate *IL6* when treated with basal conditioned media, with effects reversed by nadunolimab treatment. F) PCA plot showing fibroblasts treated with anti-IL1RAP cluster with vector control-treated fibroblasts. G) Lung metastases in oeHMGA2 tumors are significantly decreased by anti-IL1RAP treatment. H) FAP^+^ fibroblasts significantly decrease in oeHMGA2 tumors when mice receive anti-IL1RAP. αSMA^+^ fibroblasts significantly increase in oeHMGA2 tumors when mice receive anti-IL1RAP. I) Treatment with anti-IL1RAP decreases LRRC15^+^ fibroblasts and IL6^+^ fibroblasts in oeHMGA2 tumors. J) Tregs in the TME decrease in oeHMGA2 tumors when mice receive anti-IL1RAP. K) Schema for combined RASi and anti-IL1RAP treatment. L) Pancreas weights are significantly decreased by both RASi alone and RASi + anti-IL1RAP compared to control. M) Lung metastases are completely abrogated by combination therapy. N) LRRC15^+^, IL6^+^, and FAP^+^ fibroblasts are decreased by combination therapy. O) Combination therapy leads to increased CD4^+^ T cells, NK cells, and B cells in the TME. CD11b^+^ MHCII^-^ cells are significantly lower with combination therapy. C-E: Data are graphed as mean ± SD, n ≥ 2. G-I: Data are graphed as mean ± SEM, n = 10 per group. J-O: Data are graphed as mean ± SEM, n = 5 per group.

### Combining anti-IL1RAP with inhibitors of RAS

Given recent findings demonstrating the efficacy of KRAS inhibition in PDAC^53–59^, particularly in the basal subtype^14,54^, and its reliance on immunologic activity^60,61^, we next investigated RAS inhibition in combination with targeting the IL1A pathway as an approach to improve treatment efficacy in HMGA2^+^ tumors. We combined anti-IL1RAP treatment with RMC-7977, a RAS(ON) selective inhibitor that binds active RAS regardless of mutation status^56^. Syngeneic mice were implanted with 6419c5, an HMGA2^+^ T cell-low KPCY clone, and dosed with vehicle, RMC-7977, anti-IL1RAP, or both for 24 days (**Fig. 7K, Fig. S7H**). Both RMC-7977 and the combination RMC-7977 plus anti-IL1RAP led to significantly lower overall pancreas weights, with the combination trending lower (**Fig. 7L**). While all control mice (4/4), RMC-7977 mice (5/5), and anti-IL1RAP (5/5) treated mice had histologic PDAC at necropsy, a majority of the RMC-7977 + anti-IL1RAP mice (3/5) had no evidence of disease (**Fig. S8A**). Furthermore, lung metastases were completely absent in combination-treated mice (**Fig. 7M**). Combination therapy effectively decreased the area of HMGA2^+^ tumor more than either drug alone (**Fig. S7I**). In the CAF compartment, LRRC15^+^ fibroblasts were decreased only with anti-IL1RAP treatment, while FAP^+^ fibroblasts and IL6^+^ fibroblasts were more effectively ablated by combination therapy (**Fig. 7N, Fig. S7J**). aSMA^+^ and PDPN^+^ fibroblasts were decreased in the combination arm, in line with tumor regression (**Fig. S7J**). Flow cytometry studies of intratumoral immune cells were consistent with systemic anti-tumor immunity in RASi + anti-IL1RAP therapy only, with increased CD4^+^ T cells, NK cells, and B cells, and decreased MHC-II^-^ CD11b^+^ myeloid cells (**Fig. 7O, Fig. S7K**). Our results are consistent with a level of baseline resistance to RMC-7977 in T cell-low tumors, as demonstrated by Orlen et al^60^. We show that combination therapy targeting the IL1α axis allows for improved tumor control, decreased lung metastases, fibroblast remodeling, and enhanced anti-tumor immunity in HMGA2^+^ PDAC tumors.

## DISCUSSION

We define a novel mechanism by which a lineage-associated chromatin factor controls tumoral cytokine expression, leading to an immunosuppressive fibroblast phenotype in the tumor microenvironment and downregulation of anti-tumor immunity. In patient cohorts of both primary and metastatic PDAC, HMGA2 expression is associated with high levels of IL1A, increased FAP^+^ fibroblasts, increased Tregs, and decreased CD8^+^ T cells in the surrounding tumor. We demonstrate that overexpression of HMGA2 in classical subtype tumors is sufficient to induce IL1A expression, driven by its binding to the promoter region of IL1A and displacement of the linker histone H1. By utilizing immunocompetent mouse models and primary pancreatic fibroblast cultures, we find that the fibroblast skewing effects of HMGA2 can be blocked by neutralizing IL1α, deleting IL1R1, or blocking IL1RAP. Furthermore, preventing immunosuppressive fibroblast polarization equalizes anti-tumor immune surveillance between the basal and classical subtypes. These data provide an encouraging basis for blocking IL1α signaling to improve the immune response against basal subtype PDAC.

While the IL1 family of cytokines was previously shown to activate an immunosuppressive iCAF phenotype^25^, most work in the field has focused on the better-characterized IL1β^62,63^. Both genes are under the control of a common super enhancer^64^, but do not share cellular export pathways^65^. IL1α can be displayed on the membrane of some cell types, enabling direct binding to IL1R1 on adjacent cells^48^. It is ubiquitously expressed in many cell types and is more frequently described as an “alarmin,” indicating necrotic cell death^48^. IL1α can also act as a transcription factor in the nucleus^47^. These unique characteristics of IL1α raise many open questions regarding its role in PDAC tumorigenesis and progression, and further studies in the basal subtype are warranted. Previous work in PDAC has established that *IL1B* expression by tumor-associated macrophages appears to support transcriptional alterations^66^, which may partially induce *IL1A* expression in tumor cells. Autocrine IL1α signaling provides another potential source of continuous HMGA2/IL1A activation in the basal subtype. Indeed, studies in mouse models have shown that IL1α is necessary for the development of PDAC^67^. High *IL1A* expression in tumor cells has been investigated as a potential biomarker for poor prognosis in PDAC^68^. We present the intriguing finding that HMGA2 directly regulates *IL1A* expression, thus allowing HMGA2 to serve as a biomarker not only of the basal subtype^13^, but also of the associated tumor microenvironment.

We also find that IL1α can induce *LRRC15* expression in treated fibroblast cell lines and *in vivo*. These findings extend the work of Krishnamurty, et al., which showed that the LRRC15^+^ myofibroblast phenotype is conserved across cancers and is associated with a lack of response to immune checkpoint blockade and a paucity of CD8^+^ T cells^32^. These findings mirror our own and provide a potential explanation for the failure of combination immune checkpoint blockade and IL1β blockade in the clinic^63^. However, Krishnamurty and colleague attribute *LRRC15* expression on CAFs to TGFβ signaling, which remains unchanged in our oeHMGA2 model systems. The interaction between IL1 family and TGFβ signaling and how it contributes to *LRRC15* expression requires further study. IL1 and TGFβ have been described to have antagonistic roles on JAK/STAT signaling in fibroblasts^25^. Thus, more mechanistic studies are necessary to fully define the role of IL1α in regulating CAF *LRRC15* expression.

Recently, allele-specific and pan-RAS inhibitors have shown activity in clinical trials^53,56–58^. These exciting advances in targeting a previously “undruggable” oncogene have the potential to substantially improve outcomes for patients with PDAC. Safety data have been encouraging, and efficacy is currently being assessed in multiple clinical trials^53,58^. However, many resistance mechanisms have already been described, including epithelial-to-mesenchymal transition^14,69^. Preclinical studies indicate that the basal transcriptional state is especially sensitive to KRAS inhibition^14,54^, however other studies demonstrate that durable response to KRAS inhibition requires a robust T cell response^54,55,60^. Our data support the therapeutic avenue of combining IL1RAP blockade to remodel the TME alongside KRAS inhibition. Anti-IL1RAP therapies may be especially effective in other HMGA2^high^ tumors, including colorectal^70,71^, breast^72,73^, lung^74,75^, and ovarian cancers^76–78^.

Our study has several key limitations. First, there are no commercially available PDAC CAF cell lines with which to perform fibroblast skewing studies. PDAC CAFs have been described to arise from multiple different cell types, including PSCs^79,80^, bone-marrow-derived macrophages^34^, and mesenchymal cells^26,81,82^. Thus, no single cell type provides a suitable model system for studying PDAC fibroblast diversity. Furthermore, fibroblasts are highly dependent on their surrounding matrix for functionality^83,84^, and studying them without this matrix prevents a full understanding of their functionality *in vivo*. We have attempted to address these concerns by utilizing PSCs and naïve pancreatic fibroblasts, and by confirming all conclusions *in vivo*. However, given the huge variety of CAF subtypes^26,28,36,85–87^, the effect of HMGA2-driven IL1α signaling may vary widely depending on the local extracellular matrix and CAF polarization in the TME.

Second, our studies utilized orthotopic implantation as the closest proxy model for pancreatic cancer tumorigenesis. There are inherent inflammatory properties associated with direct cell injection into the pancreas, and some level of tissue damage is unavoidable. Additionally, YFP expressed by KPCY cell lines may have some immunogenic properties. While GEMMs would allow for the study of PDAC tumor initiation and progression, there are currently no conditional overexpressing HMGA2 alleles available. Thus, our studies used overexpression of LIN28B, an upstream regulator of HMGA2, to model the TME in HMGA2-driven PDAC. The reproducibility of our results between orthotopic cell line injection models and GEMM tumors is encouraging for the future translatability of this work.

Finally, targeting IL1RAP to disrupt IL1α signaling is less selective than direct IL1α neutralization or IL1R1 antagonism, as IL1RAP inhibition disrupts a wider signaling network than specific anti-IL1α antibodies or anakinra, an IL1 receptor antagonist. Both strategies have been previously studied in PDAC^49^ but have stalled in early-stage clinical trials. IL1RAP, the necessary receptor dimer protein to IL1R1, is also involved in IL-33 and IL-36 signaling. Unlike IL1R1, IL1RAP is upregulated in cancer, providing more specificity than targeting IL1α in the entire body^88^. Encouraging safety and efficacy data from clinical trials also highlights the viability of this therapeutic strategy^51^. Nadunolimab has recently been granted fast track designation by the United States Food and Drug Administration. Finally, simultaneously targeting IL-33 and IL-36 signaling in addition to IL1α signaling may have benefits in pancreatic cancer. Both cytokines have been described as having pro-cancer effects^89–91^, and blockade may allow for additional abrogation of downstream signaling from IL1α. Donahue et al. have previously described KRAS-driven IL33 as an independent mechanism of lymphocyte exclusion from the PDAC TME.^92^ While we demonstrate that IL1R1 signaling is responsible for HMGA2-driven lymphocyte exclusion, these pathways may overlap in human PDAC. Furthermore, we demonstrate that IL1RAP blockade enhances the efficacy of RASi, providing the opportunity for complete response in HMGA2-expressing PDAC.

## METHODS

### Mice

Mice were housed in pathogen-free animal facilities and experiments were conducted under protocols PROTO201900024 and PROTO201900904 in accordance with the Fred Hutchinson Cancer Center IACUC guidelines. K28C and KC mice were generated as described previously^17^ and maintained on a C57BL/6 background. IL1R1^-/-^ mice were purchased from Jackson Labs (#003245) and maintained on a C57BL/6 background. C57BL/6 (#000664) and JAXboy (#033076) mice were purchased from Jackson Labs. Data presented include both male and female mice unless noted otherwise. Experimental K28C, and KC mice were fed with 200ug/ml doxycycline in their drinking water upon weaning.

### Fibroblast isolation

Naïve pancreatic fibroblasts were derived from freshly isolated pancreata of C57BL/6 and IL1R1^-/-^ mice between 6-8 weeks of age. Tissue was minced with sterile razor blades, rinsed with DMEM:F12, then digested in 1 mg/mL collagenase II (Gibco) in DMEM:F12 for 20 minutes at 37°C and 15rpm. Digestion was halted with culture medium (DMEM:F12 supplemented with 10% FBS and 1% pen-strep) then centrifuged and resuspended in culture medium. Digested tumor tissue was passed through a 70 µm cell strainer using the plunger of a 3 mL syringe, then plated onto rat tail collagen (Corning) coated plates.

### Murine organoid and cell line isolation

Mouse pancreatic organoids were derived and cultured according to the methods detailed in Tuveson Laboratory Murine and Human Organoid Protocols (https://tuvesonlab.labsites.cshl.edu/wpcontent/uploads/sites/49/2018/06/20170523_OrganoidPro tocols.pdf). Tumor-bearing pancreatic tissue was minced with sterile razor blades and digested in digestion media [DMEM + glucose, L-glutamine, sodium pyruvate (GIBCO) with 10% fetal bovine serum and 1% penicillin (100 U/ml)/streptomycin (100 Ug/ml) (Invitrogen GIBCO)] containing dispase (0.125 mg/ml, GIBCO) and collagenase II (0.125 mg/ml, GIBCO) for 2 hours at 37°C and 15rpm. A fraction was seeded onto rat tail collagen coated plates to generate cell lines from the same tumors. The other tissue was further dissociated with TrypLE (GIBCO) for 15 minutes at 37°C and 15rpm. Cells were embedded in Matrigel and cultured in Mouse Feeding Medium (AdDMEM/F12 medium supplemented with HEPES [1×, Invitrogen], Glutamax [1×, Invitrogen], penicillin/streptomycin [1×, Invitrogen], B27 [1×, Invitrogen], Primocin [1 mg/ml, InvivoGen], N-acetyl-L-cysteine [200 μg/ml, Sigma], RSPO1-conditioned medium [10% v/v], Mouse Noggin [0.1 μg/ml, Peprotech], Mouse EGF [50 ng/ml, Peprotech], Human Gastrin [0.21 μg/ml, Tocris], Human FGF10 [0.1 μg/ml, Peprotech], Y-27632 [0.21 μg/ml, Sigma], Nicotinamide [0.045 mg/ml, Sigma], and A83-01 [0.21 μg/ml, Tocris]). To passage, murine organoids were disrupted in ice-cold splitting media (AdDMEM/F12 medium supplemented with HEPES [1×, Invitrogen], Glutamax [1×, Invitrogen], penicillin/streptomycin [1×, Invitrogen]) and re-seeded into Matrigel (Corning). Media was supplemented with 1 ug/ml doxycycline unless stated otherwise.

### Cell line xenografts

All mouse procedures were conducted in accordance with the Fred Hutchinson Cancer Center IACUC guidelines. For orthotopic injections, 3-5x10^5^ cells in 30 µL of 50% Matrigel (Corning) in DPBS were implanted into the pancreas of C57BL/6, IL1R1^-/-^, or JAXBoy mice. Briefly, anesthetized mice were prepped for surgery by removing hair and performing surgical scrubs, then a vertical incision was made in the flank to expose the peritoneum overlying the pancreas. A second small incision was made in the peritoneum, then the pancreas was herniated outside the peritoneal cavity using a sterile cotton swab until the head of the pancreas was visualized. Cells were injected, the pancreas was eased back into the abdomen with sterile cotton swabs dipped in sterile DPBS, the peritoneum was closed with adsorbable sutures (Vicryl, Ethicon Inc.), and the skin was closed with sterile wound clips (7 mm reflex, Braintree Scientific). Mice were monitored three times a week by abdominal palpation for tumor development and tumors were confirmed by ultrasound.

### Congenic bone marrow transplantation

6–8-week-old recipient mice (JAXBoy (Jackson Labs #033076) and IL1R1^-/-^ (Jackson Labs #003245)) were irradiated with 1100 rad by Cesium-137 irradiator split over 2 sessions 4 hours apart. Recipient mice were reconstituted with bone marrow from donor mice (JAXboy (Jackson Labs #033076) and IL1R1^-/-^ (Jackson Labs #003245)). Tibia, femur, and hip bones were crushed with a pestle and mortar and filtered through a 70-μm filter. Cells were subjected to RBC lysis (Thermo Fisher Scientific #00-4300-54). 10 million donor cells were retro-orbitally injected into recipient mice. Recipients were kept on Baytril water (1 mL of 100 mg/mL added to 16 oz of water) for 4 weeks and then returned to normal water for 4 weeks to allow for hematopoietic reconstitution prior to tumor implant.

### Flow cytometry

Freshly dissected tumor tissue was weighed, then minced with razor blades and digested in 10 mL of DMEM supplemented with 2480 U/mL Collagenase IV (Worthington), 1 mg/mL soybean trypsin inhibitor (Fisher), 0.5 mg/mL DNase I (Roche), and 1 mg/mL hyaluronidase (Sigma) for 45 minutes at 37°C with constant shaking at 250 rpm. Samples were vortexed every 15 minutes to aid dissociation. Digestion was quenched with 10 mL FACS buffer (DPBS (Gibco) + 3% Newborn Calf Serum (Peak Serum) + 1 mM EDTA (Teknova)), then mashed through a 100-µm cell strainer with a syringe plunger. Cells were spun down for 5 min at 500g, then counted by Vi-CELL BLU analyzer (Beckman Coulter). 2.5 x 10^6^ cells were aliquoted for staining, washed with DPBS (Gibco), and incubated with Zombie UV^TM^ (Biolegend) for 20 minutes at room temperature in the dark. Following an additional DPBS rinse, Fc blocking was performed using Fc receptor-block (anti-CD16/CD32, BioXcell, RRID:AB_2736987) on ice for 10 min. Cells were then incubated with cell surface antibody mix (see Table S1) diluted in FACS buffer and Brilliant Violet stain buffer (BD Biosciences) for 30 min on ice in the dark according to manufacturer instructions. As necessary, cells were washed twice and incubated with streptavidin fluorophores (see Table S1) for 30 min on ice in the dark. Cells were washed twice, fixed, and permeabilized with Cytofix/Cytoperm (BD Biosciences) following manufacturer instructions. Intracellular staining (see Table S1) was performed in FACS buffer and Brillian Violet stain buffer (BD Biosciences) for 30 min at room temperature in the dark. Cells were washed twice, then resuspended in FACS buffer for data acquisition. Flow cytometry was performed on a BD FACSymphony A5 flow cytometer (BD Biosciences). Data was analyzed using FlowJo (Treestar) software.

### *In vivo* anti-IL1RAP treatment

Wild-type mice with orthotopic implants of 2838 pBabe and 2838 oeHMGA2 KPCY cell lines were treated via intraperitoneal injection with 10 mg/kg murine anti-IL1RAP (clone 3A9, Cantargia) or IgG2A isotype control (MAB003, R&D Biosystems) in DPBS. Mice were dosed twice on day 1 following surgery, then twice weekly until endpoint was reached.

### *In vivo* anti-IL1RAP and RMC-7977 treatment

Wild-type mice with orthotopic implants of 6419c5 KPCY cell lines were randomized to either: 1) vehicle + mIgG2A isotype control (10 mg/kg, MAB003, R&D Biosystems); 2) RMC-7977 (10 mg/kg, #200966, ChemShuttle) + mIgG2A; 3) vehicle + anti-IL1RAP (clone 3A9, Cantargia); or 4) RMC-7977 + anti-IL1RAP. Anti-IL1RAP was dosed via intraperitoneal injection twice on day 1 following surgery, then twice weekly until endpoint. RMC-7977 (10 mg/kg, vehicle = 60% water, 20% PEG400, 10% DMSO, 10% Solutol HS15) was dosed via oral gavage on day 1 following surgery, then daily on weekdays until endpoint.

### Cell culture

PDAC cell lines HPAFII, Panc03.27, Panc1, BxPC3, AsPC1, Capan2, and CFPAC1 were obtained from ATCC (Rockville, MD). DanG was obtained from DSMZ. KP4 and SUIT2 were obtained from JCRB. Human pancreatic stellate cells (PSCs) were obtained from ScienCell (#3830) and cultured in stellate cell medium (#5301, ScienCell) to generate multiple low-passage freezes. KPCY cell lines (2838c3, 6419c5) were obtained from Kerafast (#EUP013-FP, #EUP006-FP). Panc03.27, BxPC2, AsPC1, DanG, and SUIT2 were grown in RPMI 1640 with 10% fetal bovine serum (HyClone, Cytiva). HPAFII, Panc1, Capan2, CFPAC1, human PSCs, KPCY cells, and K28C/KC murine cell lines were grown in DMEM with 10% FBS. Cells were passaged by trypsinization (0.25% Trypsin-EDTA), and studies were performed on cells cultured for fewer than twenty passages. PSCs were passaged using 0.05% Trypsin-EDTA and studies were performed on cells cultured for fewer than twelve passages.

### Constructs and viral infection

oeHMGA2 human cell lines were generated as described previously^17^. shHMGA2 knockdown human cell lines were generated using SMARTvector inducible shRNA (Clone ID #V3IHSMCB_5694017, #V3IHSMCG_8782322). oeHMGA2 murine cell lines were generated using pBabe zeo HMGA2 (gift from Tyler Jacks, Addgene plasmid #17411). shHMGA2 knockdown mouse cell lines were generated using SMARTvector inducible shRNA (Clone ID #V3IMMMCG_15449180, V3IMMMCG_16768652). Viral particles containing the above-mentioned plasmids were synthesized using either lentiviral (pCMVdR8.91) or retroviral (pCL-ECO) packaging plasmids with pCMV-VSV-G (Addgene). Cells were infected by incubating with virus and 10 µg/ml polybrene. Twenty-four hours later, cells were selected in 800 ug/mL Zeocin or 1-30 ug/ml puromycin for at least one week. Clones were produced by serial dilution in 96-well plates. For all experiments involving doxycycline-inducible HMGA2 overexpression, cells were treated with 100 ng/mL doxycycline for 72 hours. For all experiments involving doxycycline-inducible HMGA2 knockdown, cells were treated with 1 µg/mL doxycycline for 72 hours.

### Reverse siRNA knockdown

ON-TARGETplus siRNA was purchased from Dharmacon and 10 nM was reverse transfected into cells with Lipofectamine RNAiMax (Invitrogen). Non-targeting siRNA (D-001810-10-20), human *siHMGA2* SMART pool (L-013495-00-0005), mouse *siHmga2* SMART pool (L-043585-01-0010), and human *siIL1A* SMART pool (L-007952-00-0005) were used. After 16 hours medium was replaced, and cells were harvested after 72 hours. **Conditioned media experiments.** All conditioned media was produced from 1x10^6^ cells plated in 10 mL of media for 72 hours. Conditioned media was harvested and spun at 300g for 5 minutes to remove cells and particulate matter, then frozen at -80 until use. 1x10^6^ PSCs per 10 cm plate were plated in appropriate conditioned media and harvested at 72 hours by scraping in media. Conditioned media was treated with 1 ng/mL anti-IL1α (R&D Biosystems, human #MAB200-100, mouse #AF-400-SP), 20 ug/mL anti-IL1RAP (Cantargia, human Nadinolumab, mouse 15F12), 1 ug/mL anti-TGFβ (R&D Biosystems, MAB1835-100), or isotype control as indicated. For IL1α treatment, recombinant human IL-1α (PeproTech, 200-01A) was diluted in 0.1% BSA in PBS. For TGFβ treatment, recombinant human TGFβ (PeproTech, 100-21-10UG) was diluted according to manufacturer directions.

### ELISA

ELISA was performed for human and mouse IL1α using Quantikine ELISA Kits (R&D Biosystems, human IL1α #DLA50, mouse IL1α #MLA00) as per manufacturers’ instructions. 50 µL of serum or conditioned media (1:10 dilution) was loaded in duplicate.

### Whole cell lysates and western blot

Cells were scrape harvested in warm media, spun down at 300 g for 5 min, resuspended in cold PBS, and spun down again. Whole cell lysates were produced by resuspending the cell pellet in NP40 lysis buffer (2.5mM Tris-HCl pH7.5, 150mM NaCl, 1%NP40, 10% Glycerol, 2mM EDTA) supplemented with protease inhibitor cocktail (Complete EDTA-free, Roche Applied Science), 5µM TSA, 5mM sodium butyrate, 1mM DTT and phosphatase inhibitors (Phosphatase Inhibitor Cocktail Sets I, II and III, Calbiochem). Cell lysates were sonicated for 30 sec on/off for 10 min with 20% amplitude, then centrifuged at 21130 x g for 10 mins at 4°C. Protein concentration was quantified by BCA assay, and 20 µg total protein was electrophoresed on a 4-20% gradient polyacrylamide gel with SDS (Biorad) or 4-20% gradient Bis-Tris gels (GenScript) and electroblotted onto polyvinylidene difluoride membranes (Millipore) for 1-2 hours. Membranes were blocked with 5% non-fat milk in TBST for 1 hour, then probed with primary antibodies overnight. Bound proteins were detected with horseradish-peroxidase-conjugated secondary antibodies (1:5,000 Peroxidase Goat Anti-Rabbit IgG Antibody and 1:5,000 Peroxidase Horse Anti-Mouse IgG Antibody, Vector Biolaboratories PI-1000 and PI-2000, respectively) and Clarity Western ECL Max (Biorad) or Clarity Western ECL (Biorad). The following primary antibody dilutions were used: anti-HMGA2 (1:1000, Cell Signaling #5269), anti-b-actin (1:1000, Sigma Aldrich #A5316), anti-LIN28B (1:1000, Cell Signaling #4196), anti-vinculin (1:10,000, Millipore #V9131), anti-IL1α (1:1000, Cell Signaling #34819), anti-LRRC15 (1:50, Cell Signaling #50546S), anti-αSMA (1:10,000, Millipore #ABT1487).

### Real-Time RT-PCR Analysis

Total RNA was extracted using RNeasy Mini Kit (Qiagen, #74106) as described by the manufacturer. For cDNA synthesis, 1µg of total RNA was reverse-transcribed using the QuantiTect Reverse Transcription Kit (Qiagen, #205313). Real-time PCR was run in duplicate using iTaq(tm) Universal SYBR(R) Green Supermix (Biorad), following the manufacturer’s instructions, with a final volume of 12.5 µl of SYBR green reaction mix. Real-time monitoring of PCR amplification was performed using the CFX384 Real-Time system (Biorad). Data were expressed as mRNA levels normalized to the *ACTB* expression level in each sample and are represented as mean ± standard deviation between at least two independent experiments unless otherwise indicated in the figure legend. The primer sequences are listed in Table S2. Normalization was performed as necessary to standardize across biological replicates, indicated as “normalized expression” on y-axis and scaled as proportion of control condition.

### Chromatin immunoprecipitation

Cells were cross-linked with 1% formaldehyde for 10 min at room temperature and quenched on ice with 125 mM glycine for 5 min. Media was aspirated and cells were washed 3 times with ice cold PBS. Cells were lifted in lysis buffer (50 mM Tris.HCl pH 8.0, 2 mM EDTA pH 8.0, 0.1% NP40, 10% glycerol) and kept on ice for additional 5 min. Nuclei were collected by centrifugation at 300 × g for 10 min at 4 °C followed by resuspension in nuclear resuspension buffer (50 mM Tris.HCl pH 8.0, 5 mM EDTA pH 8.0, 1 % SDS). Nuclei were sonicated with Diagenode Bioruptor to an average DNA length of 150–300 bp. After centrifugation, the soluble chromatin was diluted 1:20 with DB dilution buffer (50 mM Tris.HCl pH 8.0, 5 mM EDTA pH 8.0, 0.5% NP40, 0.2 M NaCl) and 25 μg chromatin was incubated overnight with 2 µg of antibodies specific for HMGA2 (Genetex, #GTX100519), H3K4me3 (Millipore, #07-473), H3 (Abcam, #ab1791), H1 (Proteintech, #18201-1-AP), H1.5 (abcam, #ab18208), and IgG as a control (Santa Cruz Biotechnology, # sc-3877). Agarose A/G beads (Santa Cruz) added for two hours, then washed sequentially with low salt washing buffer (20 mM Tris.HCl pH 8.0, 2 mM EDTA pH 8.0, 1% NP40, 0.1% SDS, 0.15 M NaCl), high salt washing buffer (20 mM Tris.HCl pH 8.0, 2 mM EDTA pH 8.0, 1% NP40, 0.1 % SDS, 0.5 M NaCl), LiCl washing buffer (10 mM Tris.HCl pH 8.0, 1 mM EDTA pH 8.0, 1% NP40, 1% Na-deoxycholate, 0.25 M LiCl) and twice with TE buffer (10 mM Tris.HCl pH 8.0, 1 mM EDTA pH 8.0). Precipitated chromatin was reverse-crosslinked by incubating with 100 μl of 1% SDS, 0.1 M NaHCO3 and 0.5 µl RNase A (1 mg/ml) for 30 min at 37 °C. Then 2.5 µl of Proteinase K (10 mg/mL) was added for 2 h at 56 °C. Lastly, NaCl was adjusted to a final concentration of 0.2 M and mix was incubated for 16 h at 65°C. Immunoprecipitated chromatin was purified using the QIAquick PCR purification kit (Qiagen) and subjected to qPCR with primers in Table S3.

### Immunohistochemistry

Organs were harvested and fixed for two days in 10% normal buffered formalin, paraffin embedded, and sectioned (5 µm thick) by the Fred Hutchinson Cancer Center Experimental Histopathology Core (RRID:SCR_022612). Hematoxylin and eosin staining was performed following standard protocol. Immunohistochemistry was performed with standard deparaffinization and hydrogen peroxide incubation for 20 minutes, acidic antigen retrieval (10 mM citrate buffer, pH 6.0, 95°C for 20 min). Following two washes with water and two with TBST, slides were blocked for 1 hour at room temperature with 10% normal goat serum (Cell Signaling Technologies #5425). Primary antibodies (anti-HMGA2 (1:100, Cell Signaling #5269), anti-CD8α (1:200, Cell Signaling #98941), anti-LRRC15 (1:100, Cell Signaling #50546) were added to slides and incubated at 4°C overnight. The following day, slides were washed three times with TBST, then incubated with secondary antibody (biotinylated goat anti-rabbit (Vector Labs #BA-1000), biotinylated goat anti-mouse (Vector Labs #BA-9200)) for 45 minutes. Slides were probed and developed using VECTASTAIN Elite ABC Kit (Vector Labs #PK-6100) and DAB Substrate Kit (Vector Labs #SK-4100) following manufacturer’s instructions. Counterstaining was performed with hematoxylin before slides were dehydrated and mounted using Permount (Fisher).

### Immunofluorescence

Slides were sectioned, deparaffinized, and antigen retrieval was performed as described above. Following two washes with PBS, slides were permeabilized with 1x PBS/0.2% w/v gelatin/0.25% v/v Triton), then blocked with 5% normal goat serum (Cell Signaling Technologies #5425) and 1% BSA in permeabilization solution for 1 hour. Primary antibodies (anti-IL6 (1:100, abcam #ab290735), anti-podoplanin (1:200, BioLegend #127403)) in blocking buffer were added to slides and incubated at 4°C overnight. The following day, slides were washed twice with PBS and twice with permeabilization solution, then secondary antibodies were added (1:500 goat anti-Syrian hamster AlexaFluor 647 (Thermo #A-21451), 1:200 goat anti-rabbit AlexaFluor 488 (Invitrogen #A11034)) and incubated for 1 hour. Following three washes in TBST, slides were mounted using Vectastain Anti-fade Mounting Media (Vector Labs #H-1200-10).

### Multiplex immunofluorescence

Triplex IF staining was completed using a Discovery Ultra auto stainer with the following steps. Deparaffinization of the tissue followed by cell conditioning and incubation with CC1 (tris-based buffer) 48 minutes for antigen retrieval. Endogenous peroxidase activity was blocked by incubating with Discovery inhibitor (Ventana 760-4840) for 12 minutes. Slides were rinsed and FAPa (Abcam 207178) antibody was applied at a dilution of 1:50 and incubated for 40 minutes at 37C. A linking antibody of anti-Rabbit HQ (Ventana 760-4815) for detection was applied and incubated for 16 minutes at 37C followed by incubation with a Goat Ig block (Ventana 760-6008) for 4 minutes. The enzyme conjugate Anti-HQ-HRP (Ventana 760-4820) was applied and incubated for 16 minutes. The tissue was then incubated for 16 minutes with Rhodamine 6G (Ventana 760-244) for fluorescence. For the two additional antibody sequences the previous antibody was denatured by incubating the slide in CC2 (citrate buffer with a slightly acidic pH) for 8 minutes at 100C after completion of the sequence. The two additional antibody sequences followed the steps above with modifications in the incubation time of the linking antibody Anti-Rabbit-HQ at 36C for 12 minutes and the enzyme conjugate Anti-HQ-HRP for 12 minutes. The second antibody SMAa (Abcam ab5694) was applied at a dilution of 1:600 for 24 minutes at 37C, followed by incubation for 8 minutes with Discovery FAM (Ventana 760-243) for fluorescence. The third sequence was Vimentin(D21H3) (Cell Signaling 5741) at a dilution of 1:100 and incubation for 28 minutes at 37C followed by incubation with Discovery Cy5 (Ventana 760-238) for fluorescence. The slide was then incubated with QD DAPI (Ventana 760-4196) for 8 minutes as a counterstain. The slides were rinsed and a wet mount with ProLong Gold antifade media (Invitrogen P36934) was completed for tissue preservation and visualization.

### Image acquisition and quantification

Whole slide images were acquired on the Vectra Polaris Quantitative Pathology Imaging System (Akoya Biosciences, Marlborough, MA, RRID:SCR_025508). Representative stromal areas were selected and mean fluorescence intensity was calculated using Fiji.^93^ Tumor cells were excluded from analyses.

### Patient-derived bulk RNAseq

The study was approved by the Institutional Review Board of the University of Pennsylvania (IRB Protocol #830150/UPCC#13218) under a waiver of informed consent and a HIPAA waiver of authorization because it involved retrospective analysis of de-identified banked tumor specimens. Samples were retrieved from the Tumor Tissue Acquisition Bank at the University of Pennsylvania and sections were sent to Tempus for bulk RNA-seq analysis.

### Patient-derived microdissected RNAseq

RNA-seq was performed on frozen laser capture microdissected tumors on the Illumina platform. Reads were aligned using STAR (v.2.7.4a) to the human reference genome (hg38; Ensemble v100). Gene expression was calculated in transcripts per million reads mapped using stringtie (v2.0.6).

### Fibroblast RNAseq

Total RNA was isolated using the Qiagen RNeasy Kit (Qiagen) according to the manufacturer’s instructions. RNA concentration and integrity were assessed using fluorometric quantification (Nanodrop) and capillary electrophoresis (Agilent TapeStation). Samples meeting quality criteria were used for downstream library preparation. Ribosomal RNA was depleted using the Illumina Ribo-Zero Plus rRNA Depletion Kit (Illumina) following the manufacturer’s instructions. Strand-specific RNA-seq libraries were prepared using the Illumina cDNA Synthesis and RNA Prep Ligation kits, following manufacturer’s protocol. Unique dual indices were incorporated using the Illumina DNA/RNA UD Indexes Set A in combination with IDT for Illumina RNA Index Anchors to enable multiplexed sequencing. Libraries were amplified by PCR and purified using AMPure XP beads (Beckman Coulter). Library size distribution and quality were assessed using an Agilent TapeStation, and concentrations were determined by Qubit quantification. Libraries were pooled at equimolar concentrations prior to sequencing. Sequencing was conducted on an Illumina NovaSeq X Plus system (Illumina, San Diego, CA) across two lanes of a 10B-300 flow cell using a paired-end 50 bp read configuration. Base calling and demultiplexing were performed using Illumina’s standard software. Raw sequencing reads were processed using the nf-core/rnaseq pipeline (v3.22.293)^94^. Adapter trimming and quality filtering were performed prior to alignment to the mouse reference genome (GRCm39). Alignment and downstream quality control metrics were generated using default parameters of the pipeline. Median of Ratios normalization and differential gene expression analysis was performed through pyDeSeq2 v0.5.4^95^. Analyses with significantly shifted z-score distribution were corrected using fdrtool v1.2.17^96^. Top significant differentially expressed genes were identified (alpha = 0.05 and log fold change (lfc) >= 1) for gene ontology analysis through goatools v1.6.4^97^. Genes were scored by lfc * -log10(p-val) for pre-ranked gene set enrichment analysis against all collections of mouse gene sets in MSigDb v2026.1 using GSEA v4.4.0^98,99^ using the MSigDB Mouse Gene Symbol Remapping v2026.1.Mm Chip file with the Collapse option. Additional gene sets were identified from^28,29,36^ for analysis. Comparison of gene expressions were evaluated using the t-test implemented in scipy^100^. All plots were constructed using matplotlib v3.10.8 and decoupler-py v2.1.5^101^.

### Spatial transcriptomics analysis

The unique molecular identifier count matrices were downloaded from GEO (GSE274557) and analyzed using scVerse (squidpy v1.7.0^102^ and scanpy v1.11.5^103^). Spots were filtered for complexity (UMI count > 100) and for tissue inclusion. “Neighborhood counts” were generated for each spot by summing up each spot’s expression with its 6 immediate neighboring spots. To preserve spatial fluctuations in gene expression and counts, these neighborhood counts were log1p normalized to obtain the “log(1 + Neighborhood Count)” values. Samples were further grouped by tissue origin (Primary, Lung, Liver, and Peritoneal) as delineated in Supplementary Table 5 of ^35^. The iStar scores were obtained from the authors of ^35^. Spot lineage was determined by first normalizing the spot UMI counts by scanpy’s normalize_total followed by the log1p function to obtain the “normalized log1p” counts. These values were used to score each spot for Basal and Classical scores using the score_genes function with parameters n_bins=24, ctrl_size=100, and gene_list coinciding with Supplementary Table 4 of ^35^. The difference between these scores with a threshold of 0.3 and - 0.3 labeled each spot as Basal or Classical, respectively. Comparison between gene expressions and scores were evaluated using the Mann-Whitney U-test implemented in scipy^100^. All plots were constructed using matplotlib v3.10.8 and decoupler-py v2.1.5^101^.

### Statistical analysis

Statistical tests were performed with GraphPad Prism v.10.3 (RRID:SCR_002798). Statistical significance was set at p=0.05. Comparisons between two groups were analyzed with the Mann-Whitney U Test (unpaired) or Student’s t-test (unpaired).

### Data and materials availability

Generated cell lines are available by request from the corresponding author. Fibroblast RNAseq is uploaded to GEO (accession number TBD). Patient-derived bulk RNAseq data (n=79) is available at GEO. Patient-derived microdissected RNAseq data is available from F.N. upon request. KPCY RNAseq is available from GEO (accession number GSE189925).

## Acknowledgments

The authors sincerely thank Deby Kumasaka of the Holland Lab for her assistance with multiplex immunofluorescence and the Experimental Histopathology Shared Resource (RRID:SCR_022612) and the Comparative Medicine Shared Resource (RRID: SCR_022610) at Fred Hutchinson Cancer Center (P30 CA015704) for their crucial support of this study. We thank E.A. Collisson and past and present members of the Kugel Lab for their helpful discussions.

## Author contributions

N.Y., S.D, T.M.R., A.W.P, N.C., A.O, and J.A.F., collected data. N.Y., S.D., T.M.R, A.W.P., N.C., A.O., J.H.S., J.A.F., and K.C.B. analyzed data. L.D.B, A.S., and S.A. provided experimental support and performed preliminary experiments. I.L. and G.H.J. performed independent dataset analyses. G.L.B., S.R.H, C.R.M., N.G.S, D.B.V-C., R.F., H.C.C., and K.C.B. provided materials and supplementary data. N.Y., D.B.V-C., R.F., K.C.B, and S.K. contributed to experimental design. N.Y. wrote the paper. All authors discussed the results and commented on the manuscript.

## Tables

**Supplementary Table 1.** Flow antibodies.

**Supplementary Table 2.** RT-qPCR primers.

**Supplementary Table 3.** ChIP-qPCR primers.

---

**Supplemental Figure S1.**
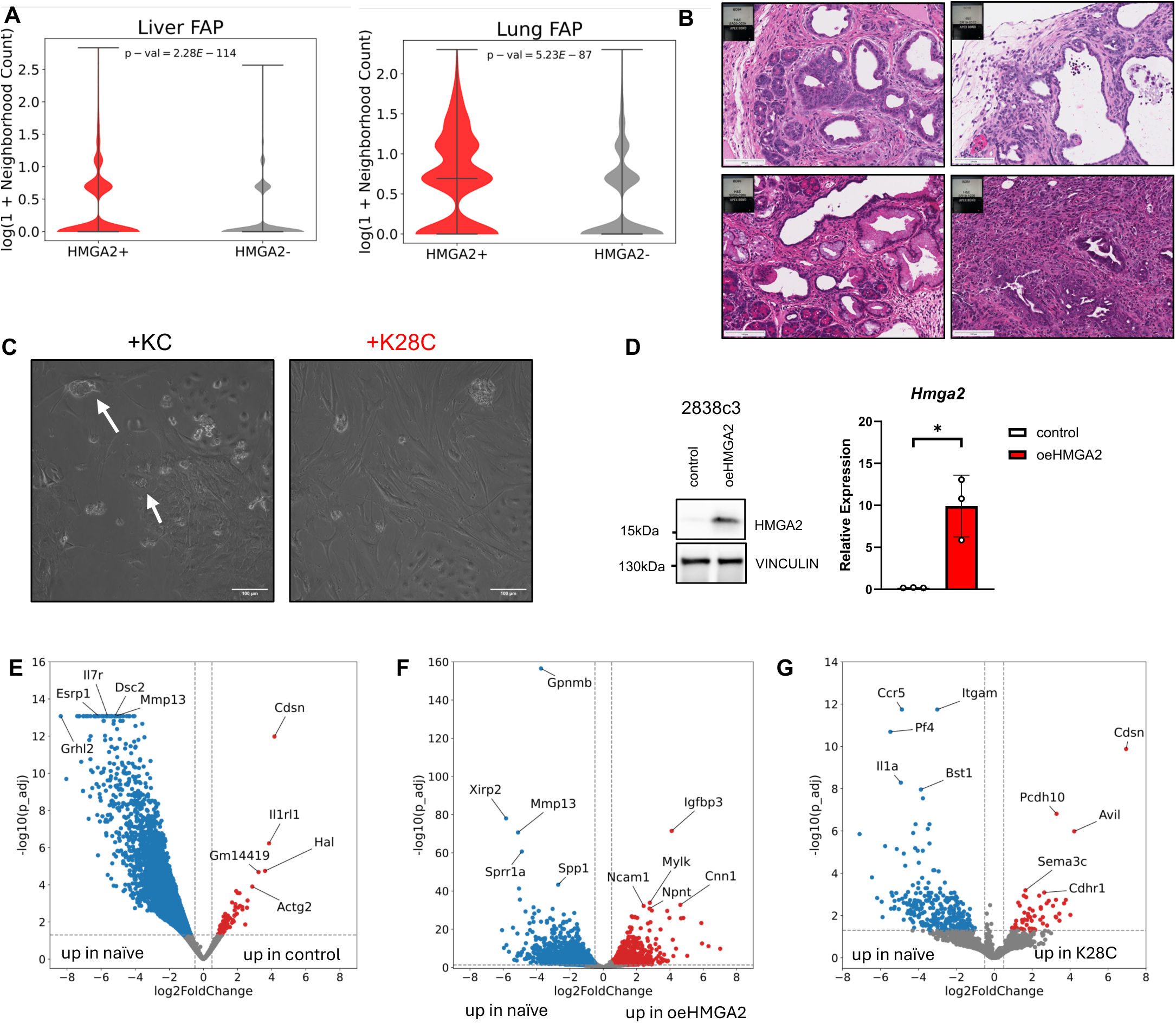
A) Metastatic tumor sites show higher *FAP* expression near basal *HMGA2*^+^ tumors compared to classical *HMGA*2^-^ tumors. B) Paired H&E micrographs for multiplex immunofluorescence in Figure 1E. Scale bar = 100 um. C) Brightfield microscopy reveals quiescent fibroblasts with lipid inclusions (arrowheads) in classical KC organoid-treated murine fibroblasts, while basal K28C organoid-treated fibroblasts take on proliferative phenotypes. Scale bar = 100 um. D) Western blot (left) and RT-qPCR validation (right) of stable *Hmga2* overexpression in the 2838c3 T cell-high KPCY cell line. E) Volcano plot of genes upregulated in KPCY vector control-treated fibroblasts versus naïve fibroblasts. F) Volcano plot of genes upregulated in KPCY oeHMGA2-treated fibroblasts versus naïve fibroblasts. G) Volcano plot of genes upregulated in K28C-treated fibroblasts versus naïve fibroblasts. Data are graphed as mean ± SD, n ≥ 3 biological replicates unless otherwise indicated.

**Supplemental Figure S2.**
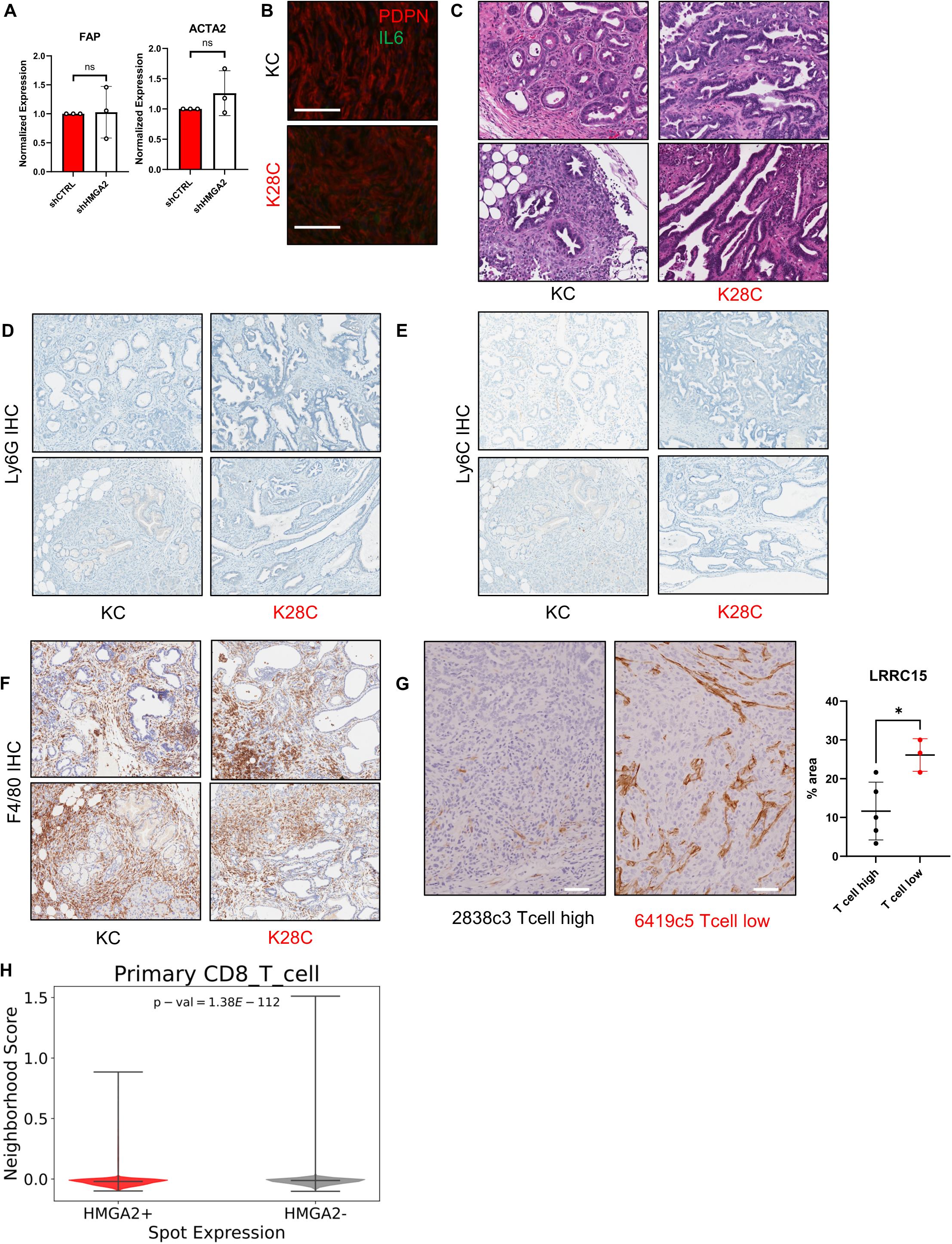
A) shRNA-mediated knockdown of *HMGA2* in three basal cell lines does not alter *FAP* and *ACTA2* expression in treated PSCs (n=3, Panc1, KP4, Panc3.27). B) Representative IL6 (green) and PDPN (red) immunofluorescence on K28C and KC tumors. Scale bar = 50 µm. C) Paired H&E micrographs for CD8 IHC from Figure 1G. Scale bar = 100 um. D) Representative Ly6G IHC from K28C and KC tumors. E) Representative Ly6G IHC from K28C and KC tumors. F) Representative F4/80 IHC from K28C and KC tumors. G) Representative images and scoring for LRRC15 immunohistochemistry on 2838c3 T cell-high and 6419c5 T cell-low KPCY cell lines injected orthotopically. Scale bar = 50 um. H) CD8 T cell signatures are higher near *HMGA2^-^* tumor spots compared to *HMGA2*^+^ spots in spatial transcriptomics.

**Supplemental Figure S3.**
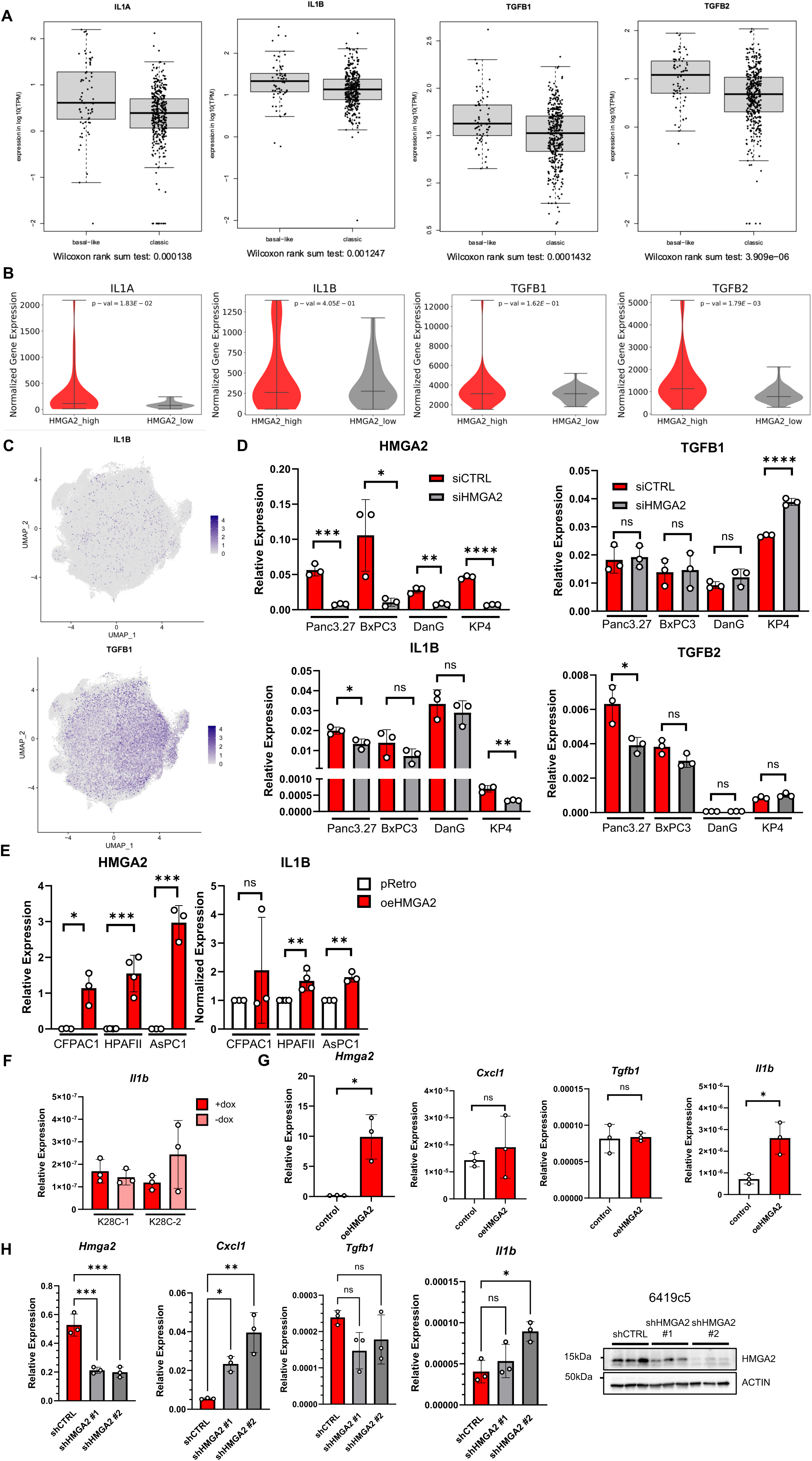
A) Microdissected RNAseq on resected tumor samples from the COMPASS trial shows higher *IL1A*, *IL1B*, *TGFB1*, and *TGFB2* expression in HMGA2^+^ samples. B) Bulk RNAseq on 79 patient tumors shows association of *IL1A* but not *IL1B, TGFB1*, or *TGFB2* with *HMGA2* expression. C) scRNAseq UMAP of ductal cell object with expression of *IL1B* and *TGFB1* expression overlaid. D) RT-qPCR validation of HMGA2 knockdown in siHMGA2 cell lines and associated *IL1B, TGFB1,* and *TGFB2* expression. E) RT-qPCR validation of HMGA2 overexpression in classical cell lines and associated *IL1B* and *TGFB1* expression. F) *Il1b* expression is unaltered in K28C mouse cell lines with dox withdrawal. G) RT-qPCR validation of *Hmga2* overexpression in 2838c3 KPCY cell line with paired *Cxcl1*, *Tgfb1*, and *Il1b* expression. H) RT-qPCR validation of *Hmga2* knockdown in 6419c5 KPCY cell line with paired *Cxcl1*, *Tgfb1*, and *Il1b* expression. Western blot validation of HMGA2 loss in shHMGA2 6419c5 cell lines. Data are graphed as mean ± SD, n ≥ 3 biological replicates unless otherwise indicated.

**Supplemental Figure S4.**
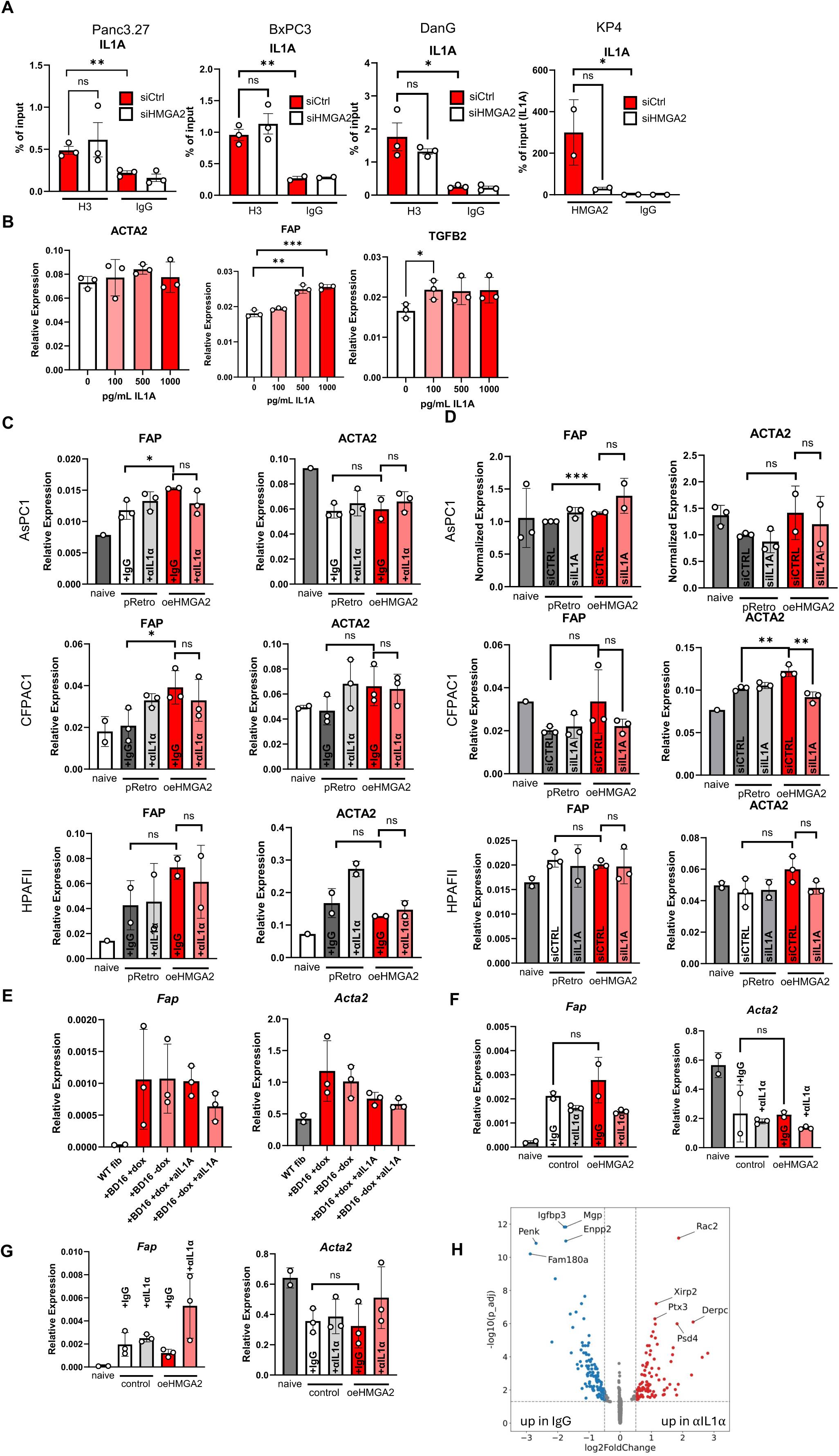
A) H3 occupancy does not change at the IL1A promoter with HMGA2 knockdown. B) *FAP* expression in PSCs increases with increasing doses of IL1α, but *ACTA2* expression is unchanged. C) Conditioned media from classical cell lines with and without overexpression of HMGA2 does not reliably alter *FAP* or *ACTA2* expression. Treatment with αIL1α has no effect. D) Conditioned media from classical cell lines with and without overexpression of HMGA2 does not reliably alter *FAP* or *ACTA2* expression. Knocking down *IL1A* in tumor cells has no effect. E) Paired *Fap* and *Acta2* expression in naïve murine fibroblasts treated with K28C conditioned media with and without αIL1A. F) Paired *Fap* and *Acta2* expression in naïve murine fibroblasts treated with 2838c3 vector control and oeHMGA2 conditioned media with and without αIL1A. G) Paired *Fap* and *Acta2* expression in naïve murine fibroblasts derived from *Il1r1*^-/-^ mice treated with 2838c3 vector control and oeHMGA2 conditioned media with and without αIL1A. H) Volcano plot of differentially expressed genes in oeHMGA2-treated fibroblasts when treated with antiIL1α versus isotype control. Data are graphed as mean ± SD, n ≥ 2 biological replicates unless otherwise indicated.

**Supplemental Figure S5.**
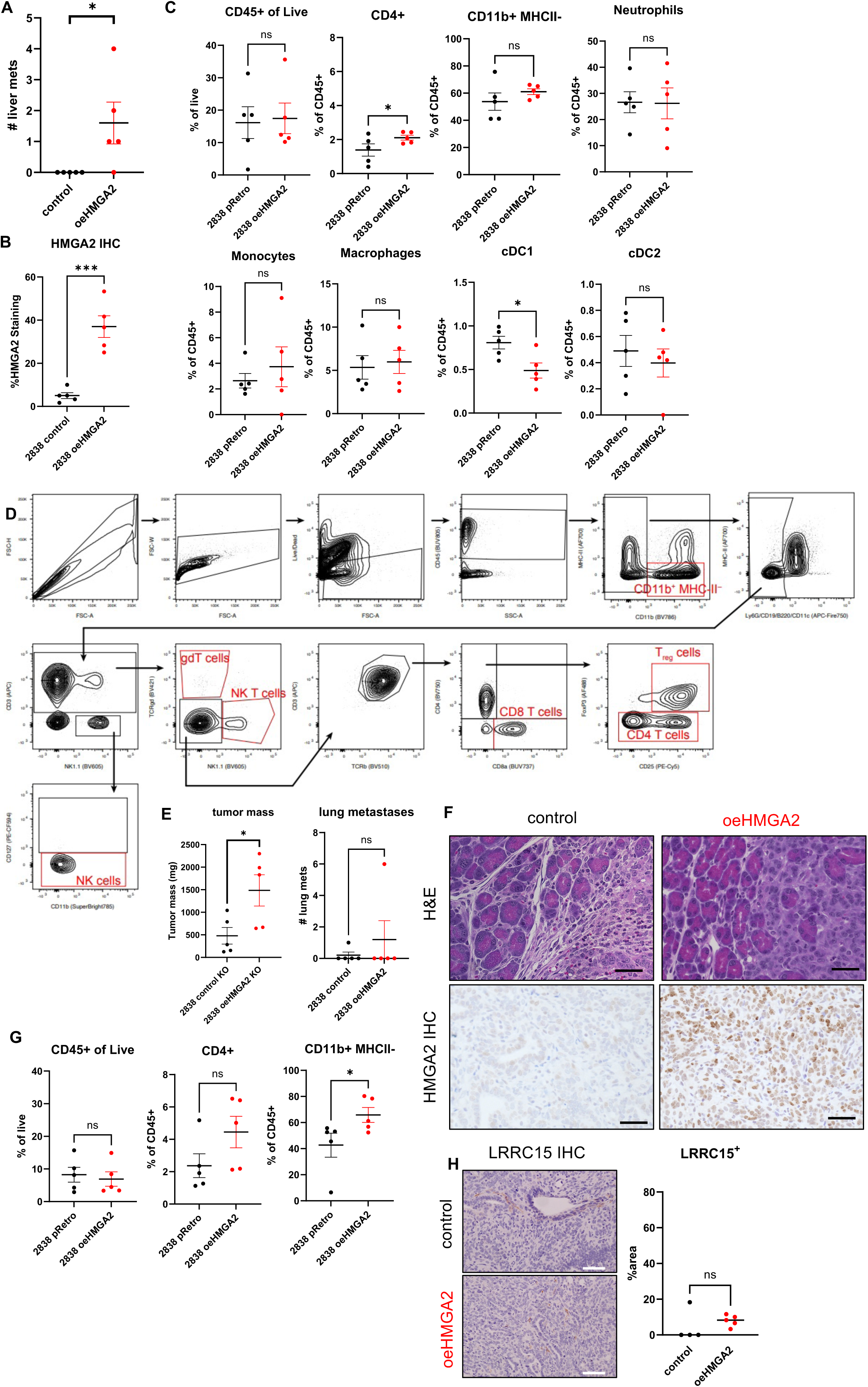
A) Liver metastases are higher in oeHMGA2 tumors compared to control tumors. B) Scoring confirms higher expression of HMGA2 in oeHMGA2 tumors. C) No significant differences are seen in total CD45^+^ infiltrate, CD11b^+^ MHC-II^-^ myeloid lineage, neutrophils, monocytes, macrophages, or cDC2s between control and oeHMGA2 tumors. CD4^+^ T cells are higher in oeHMGA2 tumors, while cDC1s are higher in control tumors. D) Flow gating scheme. E) Tumor mass is higher in oeHMGA2 tumors when implanted into *Il1r1^-/-^* hosts, but lung metastases are not significantly different. F) Representative H&E and HMGA2 immunohistochemistry of control and oeHMGA2 tumors injected into the pancreata of *Il1r1*^-/-^ mice. G) No significant differences are seen in total CD45^+^ infiltrate or CD4^+^ T cell content when *Il1r1*^-/-^ mice are used as recipients. CD11b^+^ MHC-II^-^ myeloid lineage cells are higher in oeHMGA2 tumors. H) Representative LRRC15 IHC and scoring on tumors from *Il1r1*^-/-^ mice show no differences. Data are graphed as mean ± SEM, n = 5 per group.

**Supplemental Figure S6.**
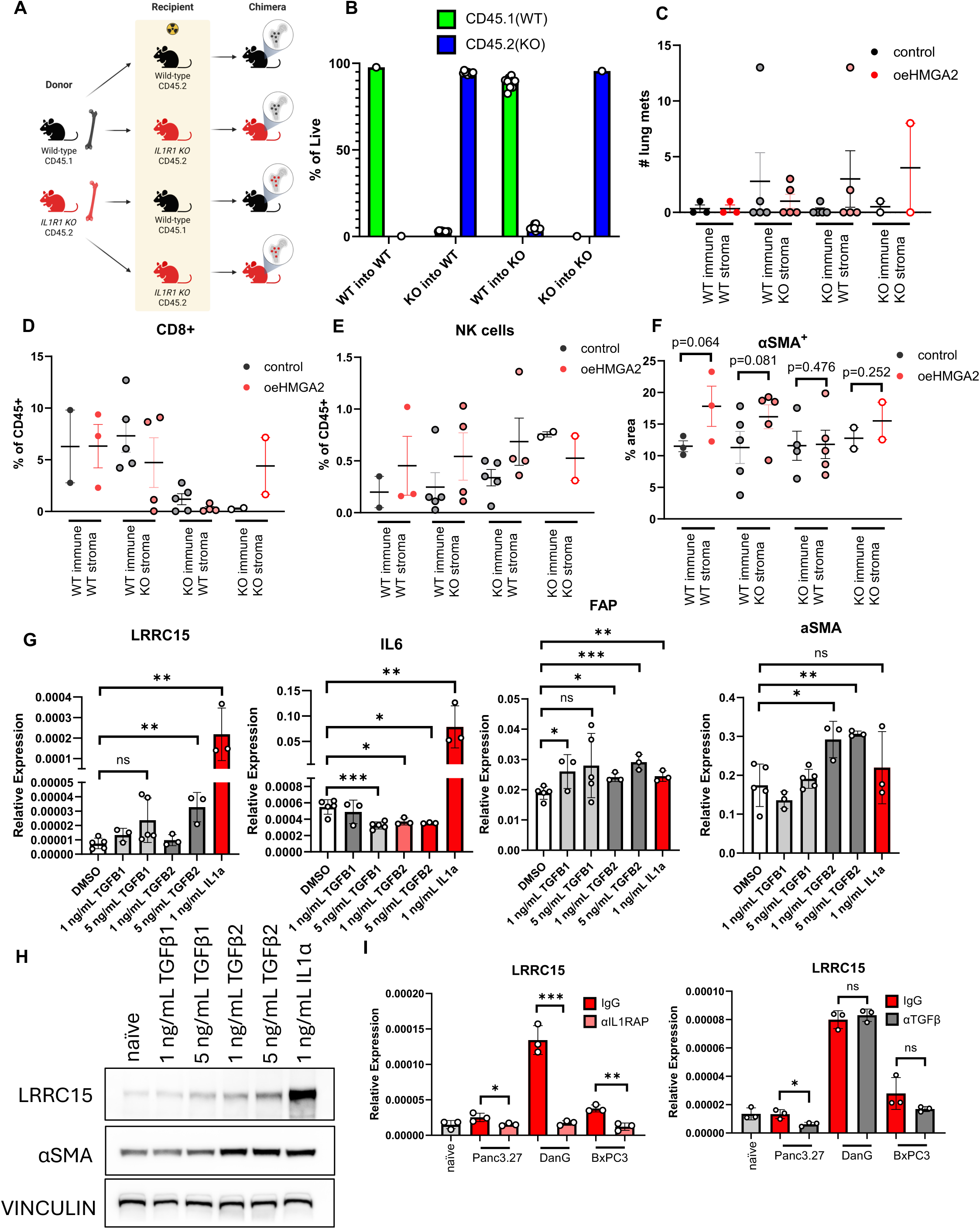
A) Expanded bone marrow chimera schema including control animals. B) Flow cytometry on peripheral blood confirms presence of CD45.1 IL1R1 WT immune cells in WT bone marrow recipients. CD45.2 IL1R1 KO immune cells are present in KO bone marrow recipients. C) Lung metastases are not significantly altered by tumoral HMGA2 expression in either chimera. D-E) IL1R1 KO CD8^+^ T and NK cells rely on effective IL1R1 signaling on both stroma and immune cells to result in differences between control and oeHMGA2 tumors. F) αSMA^+^ fibroblasts trend higher in oeHMGA2 tumors when immune cells are IL1R1 WT. G) RT-qPCR of PSCs treated with recombinant IL1α, TGFβ1, or TGFβ2 at indicated doses leads to varying upregulation of *LRRC15, IL6, FAP,* and *αSMA*. H) Western blot of PSCs treated with recombinant IL1α, TGFβ1, or TGFβ2 at indicated doses shows largest protein upregulation of LRRC15 occurs with IL1α treatment. I) Blockade of IL1RAP leads to downregulation of *LRRC15* in basal media-treated PSCs. Neutralization of TGFβ1-3 does not consistently lead to downregulation of LRRC15. A-F: Data are graphed as mean ± SEM, n ≥ 2 per group. G-H: Data are graphed as mean ± SD, n ≥ 3 biological replicates unless otherwise indicated.

**Supplemental Figure S7.**
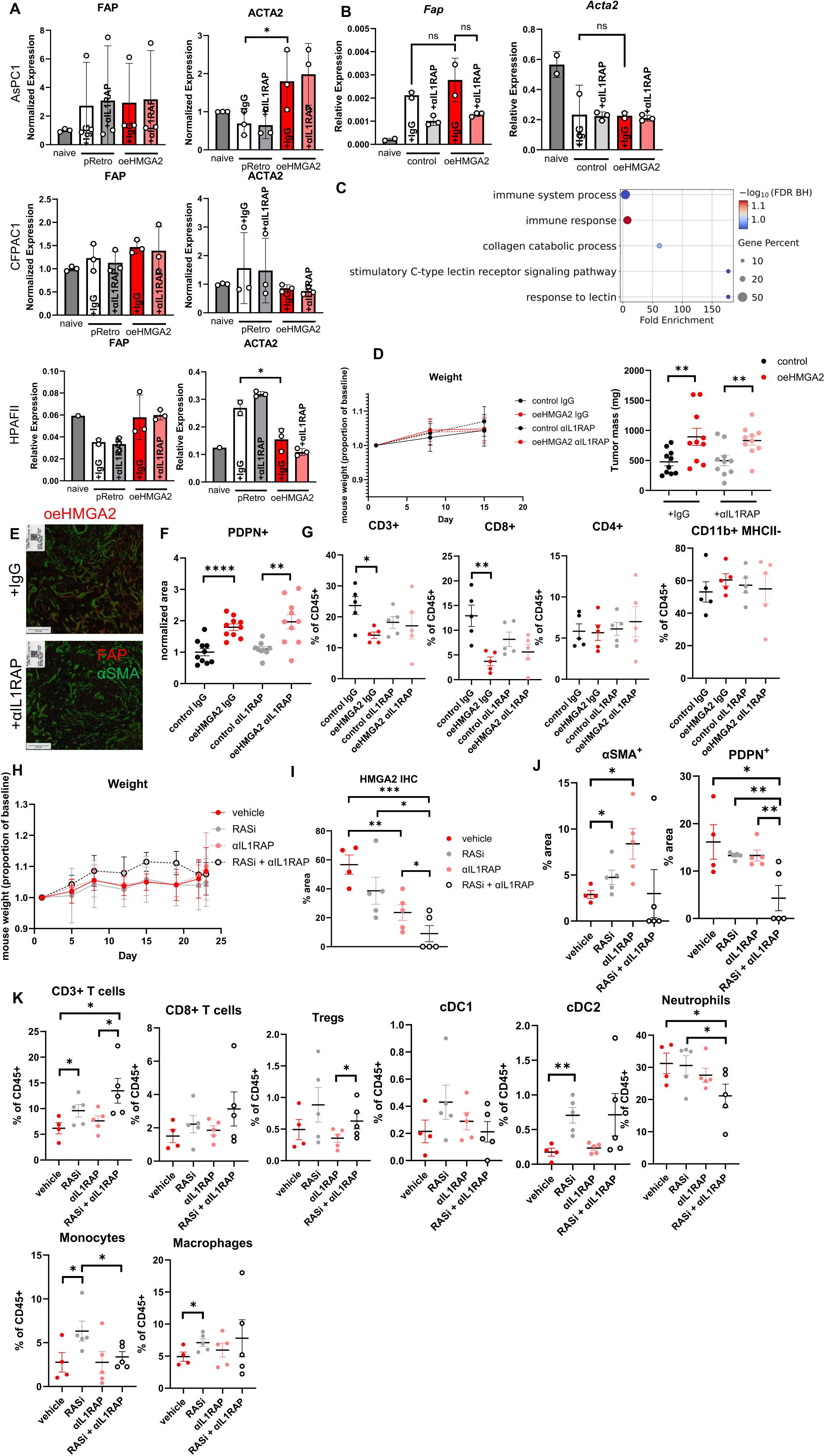
A) Paired FAP and ACTA2 expression shows minimal changes when PSCs are treated with conditioned media from indicated cell lines with or without nadunolimab. B) Paired Fap and Acta2 expression shows minimal changes when murine pancreatic fibroblasts are treated with indicated conditioned media with or without anti-IL1RAP. C) Top five GO terms enriched in oeHMGA2-conditioned fibroblasts when treated with anti-IL1RAP. D) Anti-IL1RAP treatment is well tolerated *in vivo* (left), and final tumor mass is higher in oeHMGA2 tumors in anti-IL1RAP and IgG-treated tumors (right). E) Representative dual IF for FAP and αSMA in oeHMGA2 tumors with IgG and anti-IL1RAP. Scale bar = 100 um. F) PDPN^+^ expression shows minimal effect of anti-IL1RAP treatment. G) Treatment with anti-IL1RAP does not alter CD3^+^ T cells, CD8^+^ T cells, CD4^+^ T cells, or CD11b^+^ MHCII^-^ myeloid cells in oeHMGA2 tumors. H) RASi and anti-IL1RAP is well tolerated by mice. I) HMGA2^+^ tumor area is significantly decreased by combination therapy. J) αSMA^+^ and PDPN^+^ fibroblasts scoring for combination therapy. K) CD3+ T cell, CD8+ T cell, Treg, cDC1, cDC2, neutrophil, monocyte, and macrophage populations in combination treatment TME. A-B: Data are graphed as mean ± SD, n ≥ 2. C-F: Data are graphed as mean ± SEM, n = 10 per group. G-K: Data are graphed as mean ± SEM, n = 5 per group.

**Supplemental Figure S8.**
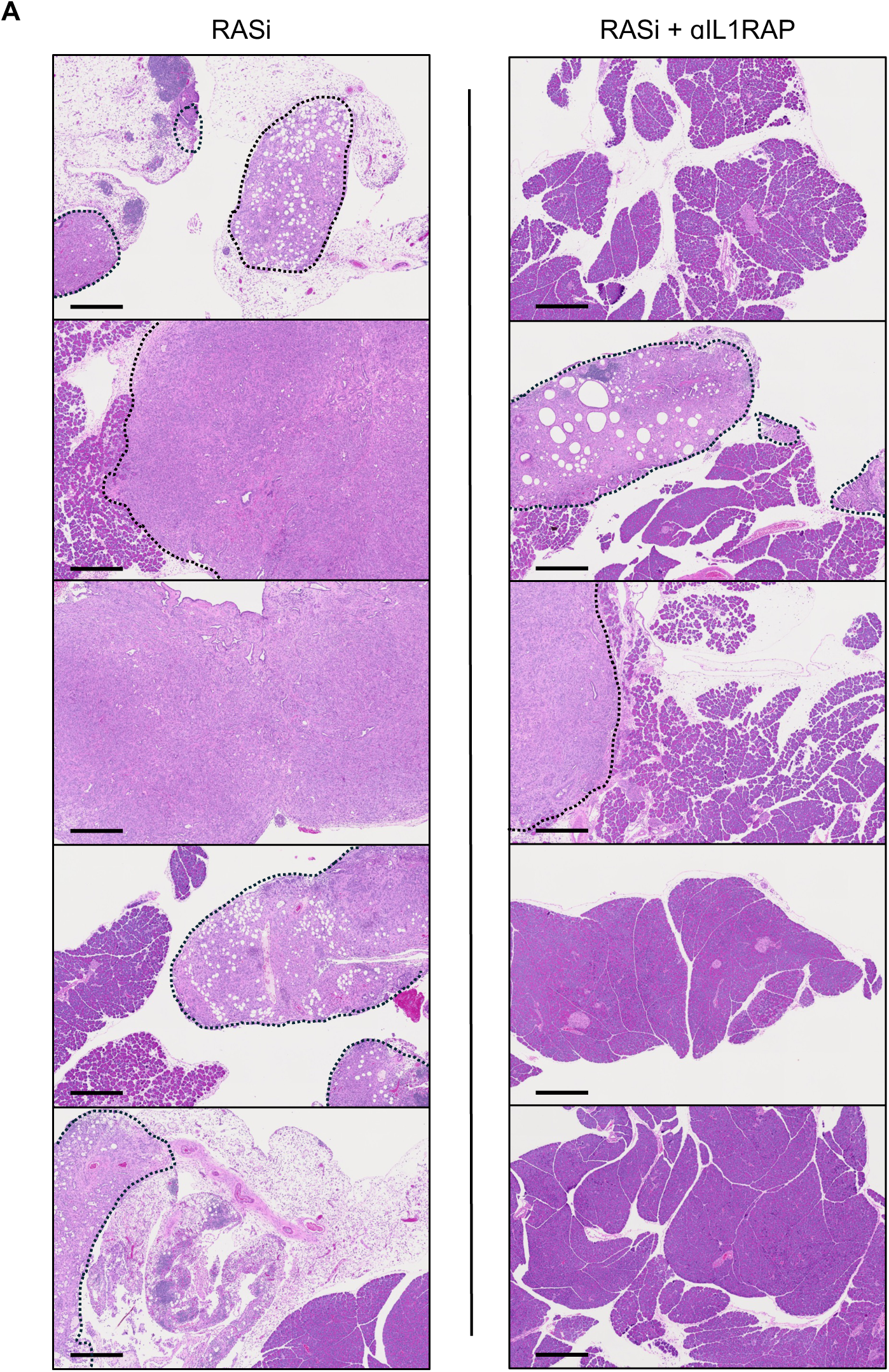
A) H&E of HMGA2^+^ tumors treated with RASi (left) or RASi and anti-IL1RAP at endpoint. Scale bar = 500 µm.

